# Antidepressant and antipsychotic drugs reduce viral infection by SARS-CoV-2 and fluoxetine show antiviral activity against the novel variants *in vitro*

**DOI:** 10.1101/2021.03.22.436379

**Authors:** Senem Merve Fred, Suvi Kuivanen, Hasan Ugurlu, Plinio Cabrera Casarotto, Lev Levanov, Kalle Saksela, Olli Vapalahti, Eero Castrén

**Affiliations:** Neuroscience Center - HiLIFE, University of Helsinki, Finland; Department of Virology, University of Helsinki, Finland; Department of Veterinary Biosciences, University of Helsinki, Finland; HUS Diagnostic Center, HUSLAB, Clinical Microbiology, University of Helsinki and Helsinki University Hospital, Finland

## Abstract

**Background and Purpose:** Repurposing of currently available drugs is a valuable strategy to tackle the consequences of COVID-19. Recently, several studies have investigated the effect of psychoactive drugs on SARS-CoV-2 in cell culture models as well as in clinical practice. Our aim was to expand these studies and test some of these compounds against newly emerged variants.

**Experimental Approach:** Several antidepressant drugs and antipsychotic drugs with different primary mechanisms of action were tested in ACE2/TMPRSS2-expressing human embryonic kidney cells against the infection by SARS-CoV-2 spike protein-dependent pseudoviruses. Some of these compounds were also tested in human lung epithelial cell line, Calu-1, against the first wave (B.1) lineage of SARS-CoV-2 and the variants of concern, B.1.1.7 and B.1.351.

**Key Results:** Several clinically used antidepressants, including fluoxetine, citalopram, reboxetine, imipramine, as well as antipsychotic compounds chlorpromazine, flupenthixol, and pimozide inhibited the infection by pseudotyped viruses with minimal effects on cell viability. The antiviral action of several of these drugs was verified in Calu-1 cells against the (B.1) lineage of SARS-CoV-2. By contrast, the anticonvulsant carbamazepine, and novel antidepressants ketamine and its derivatives as well as MAO and phosphodiesterase inhibitors phenelzine and rolipram, respectively, showed no activity in the pseudovirus model. Furthermore, fluoxetine remained effective against pseudo viruses with N501Y, K417N, and E484K spike mutations, and the VoC-1 (B.1.1.7) and VoC-2 (B.1.351) variants of SARS-CoV-2.

**Conclusion and Implications:** Our study confirms previous data and extends information on the repurposing of these drugs to counteract SARS-CoV-2 infection including different variants of concern.

## Introduction

Coronaviruses, members of the enveloped RNA virus family *Coronaviridae* (Lai & Cavanagh, 1997), are known to infect multiple species ranging from birds to mammals (To et al., 2013). In humans, in addition to four coronaviruses causing common colds, high level of pathogenicity of viruses from this family, such as severe acute respiratory syndrome coronavirus (SARS-CoV) and Middle East respiratory syndrome coronavirus (MERS-CoV), were observed in epidemics that emerged in 2003 and 2012, respectively (Q. Tang et al., 2015). The pandemic that started in December 2019 in Wuhan, China, caused by SARS-CoV-2 infection, and the dramatic increase in the number of infected people and death due to COVID-19, has shifted the efforts of scientists towards investigating this virus more closely (F. Wu et al., 2020; Zhou et al., 2020). Autopsies of patients who died from COVID-19 have reported the spread of virus to lungs, kidneys, liver, heart, brain, and blood (Puelles et al., 2020). Taken together, severity of cases and high infectivity of the virus requires more attention and effort towards controlling the global epidemic and developing treatment options.

The spike protein (S protein) localized to the viral envelope has been annotated as one of the critical parts of this group of viruses because of its role in attachment and fusion to host target cells (F. Li et al., 2005). Angiotensin-converting enzyme 2 (ACE2) located on the surface of the target cells is recognized by S protein of SARS-CoV and SARS-CoV-2 (Hoffmann, Kleine-Weber, Schroeder, et al., 2020; W. Li et al., 2003; Walls et al., 2020; Q. Wang et al., 2020). A distinct location, the receptor-binding domain (RBD), in S protein is important for binding to ACE2 (F. Li et al., 2005; Wan et al., 2020). ACE2-binding alone may allow the viral entry to target cells, however, proteolytic processing of S protein by transmembrane protease/serine subfamily member 2 (TMPRSS2) has been shown to enhance viral entry (Hoffmann, Kleine-Weber, Schroeder, et al., 2020; Shulla et al., 2011). Recent data on SARS-CoV-2 also suggests that the S protein cleavage by furin proprotein convertase (Hoffmann, Kleine-Weber, & Pöhlmann, 2020; Johnson et al., 2021) and binding of the C-end of the S1 to the cellular receptor neuropilin-1 can also augment infectivity of this virus (Cantuti-Castelvetri et al., 2020; Daly et al., 2020). Lysosomal enzyme Cathepsin L has been shown to regulate priming of SARS-CoV-2 S protein in order to provide entry of viral RNA genome into the host cytoplasm (Ou et al., 2020). Although many key players facilitating viral entry and the routes of infection have been identified since the emergence of COVID-19, effective drugs that can alleviate or eliminate the viral infection remain to be found.

Recently, a number of SARS-CoV-2 variants of concern (VoCs) have been identified including VoC-1 (B.1.1.7 lineage) first identified in the UK (*Preliminary Genomic Characterisation of an Emergent SARS-CoV-2 Lineage in the UK Defined by a Novel Set of Spike Mutations - SARS-CoV-2 Coronavirus / NCoV-2019 Genomic Epidemiology*, 2020), VoC-2 first identified in South Africa (B.1.351 lineage) (Tegally et al., 2020), and the VoC-3 originating in Brazil (B.1.1.28 lineage and the descendent P1 lineage) (*Genomic Characterisation of an Emergent SARS-CoV-2 Lineage in Manaus*, 2021; *Phylogenetic Relationship of SARS-CoV-2 Sequences from Amazonas with Emerging Brazilian Variants Harboring Mutations E484K and N501Y in the Spike Protein - SARS-CoV-2 Coronavirus /NCoV-2019 Genomic Epidemiology*, 2021; Voloch et al., 2020). Critical mutations in the RBD of S protein have been detected in these variants which can affect the ACE2 affinity (associated with N501Y shared by all three VoCs) and antibody neutralization (associated with E484K shared by VoC-2 and 3 and present in some strains of VoC-1) (*Genomic Characterisation of an Emergent SARS-CoV-2 Lineage in Manaus*, 2021; Greaney, Loes, et al., 2021; Greaney, Starr, et al., 2021; Gu et al., 2020; Nelson et al., 2021; Starr et al., 2020; Tegally et al., 2020; Wibmer et al., 2021). The emergence of particularly VoC-2 and 3 variants has put the currently available vaccines and vaccine candidates under dispute (Fontanet et al., 2021).

The strategy of drug repurposing has been used as a method of searching novel treatments for virus-related diseases (Dyall et al., 2014; Mercorelli et al., 2018). A pre-clinical study has shown that the antidepressant drugs, such as sertraline, paroxetine, and clomipramine, can reduce the *Zaire* Ebola virus (EBOV) entry to target cells (Johansen et al., 2015). In line with this, the treatment of COVID-19 patients with fluoxetine, escitalopram, and venlafaxine for 20 days has been found to reduce risk of intubation or death by COVID-19 (Hoertel et al., 2021). Another study has shown that the treatment of COVID-19 patients with fluvoxamine for 2 weeks was effective to decrease the development of clinical deterioration (Lenze et al., 2020).

In the present study, we addressed if the psychoactive drugs can be used to reduce SARS-CoV-2 infection of host cells *in vitro*. We show that pharmacologically diverse antidepressant drugs, as well as several antipsychotics were able to reduce the infection by pseudotyped viruses harboring SARS-CoV-2 S protein. Treatment of human lung epithelial cell line Calu-1 infected with the B.1 lineage of SARS-CoV-2 with these drugs was also successful in reducing the amount of infectious virus. Moreover, infection by pseudotyped viruses carrying N501Y, K417N, or E484K single point mutations or triple mutant (N501Y/K417N/E484K) in the spike protein was shown to be reduced by fluoxetine. Fluoxetine was also effective against the variants of SARS-CoV-2, VoC-1 (B.1.1.7) and VoC-2 (B.1.351), in Calu-1 cells.

## Materials and Methods

### Drugs

Fluoxetine (#H6995, Bosche Scientific), citalopram (#C505000, Toronto Research Chemicals), paroxetine (#2141, Tocris), fluvoxamine (#1033), venlafaxine (#2917, Tocris), reboxetine (#1982, Tocris), imipramine (#I7379-5G, Sigma–Aldrich), clomipramine (#C7291, Sigma–Aldrich), desipramine (#3067, Tocris), phenelzine (#P6777, Sigma–Aldrich), rolipram (#R6520, Sigma–Aldrich), ketamine (#3131, Tocris), 2R, 6R-Hydroxynorketamine and 2S, 6S-Hydroxynorketamine (#6094 and #6095, respectively, Tocris), carbamazepine (#4098, Tocris), chlorpromazine (#C8138, Sigma–Aldrich), flupenthixol (#4057, Tocris), and pimozide (#0937, Tocris) were investigated in this study. The compounds were dissolverd in dimethyl sulfoxide (DMSO) that was also used as a vehicle in all the experiments. Concentrations tested (Table 1) were selected based on earlier *in vitro* studies from our research group (Casarotto et al., 2021; Fred et al., 2019) and other laboratories (Johansen et al., 2015).

**Table 1.** Psychoactive drugs implicated in our study and the doses used for the treatment of HEK 293T-ACE2-TMPRSS2 and Calu-1 cells.

| Category of drugs tested | Name of drugs | Dose (μM) |
| --- | --- | --- |
| Selective serotonin reuptake inhibitors (SSRIs) | Fluoxetine | 0.01 - 20 |
|  | Citalopram | 0.01 - 50 |
|  | Paroxetine | 0.01 - 20 |
|  | Fluvoxamine | 0.01 - 50 |
| Selective norepinephrine reuptake inhibitor (NRI) | Reboxetine | 0.1 - 50 |
| Serotonin - norepinephrine reuptake inhibitor (SNRI) | Venlafaxine | 0.01 - 50 |
| Tricyclic antidepressants | Clomipramine | 0.01 - 10 |
|  | Imipramine | 0.01 - 50 |
|  | Desipramine | 0.01 - 20 |
| Rapid-acting antidepressants | Ketamine | 0.01 - 50 |
|  | 2R, 6R-Hydroxynorketamine (2R, 6R-HNK) |  |
|  | 2S, 6S-Hydroxynorketamine (2S, 6S-HNK) |  |
| Phosphodiesterase-4 inhibitor | Rolipram | 0.01 - 50 |
| Monoamine oxidase inhibitor | Phenelzine | 0.01 - 50 |
| Antipsychotic | Chlorpromazine | 0.01 - 5 |
|  | Flupenthixol | 0.01 - 10 |
|  | Pimozide | 1 - 10 |
| Anticonvulsant | Carbamazepine | 0.1 - 50 |

### Cell lines

TMPRSS2 (NM_001135099.1) and ACE2 (AB046569.1) cDNAs were sequentially transfected into HEK293T cells using lentiviral vectors. The positive colonies were selected by blasticidin or puromycin resistance for TMPRSS2 and ACE2, respectively. The viral vectors were generated by transfecting HEK293T cells using polyethylenimine with the packaging vector pCMV-dR8.91 and the vesicular stomatitis virus (VSV-G) envelope expression vector pMD2.G (#12259, Addgene) with either pLenti6.3/V5-DEST-TMPRSS2 (from UH Biomedicum Functional Genomic Unit) or pWPI-puro [modified from, #12254, Addgene, (Z. Zhao et al., 2019)] expressing ACE2 (cDNA from Dr. Markku Varjosalo). Six hours after transfection, media was replaced with fresh Dulbecco’s Modified Eagle Medium (DMEM) containing high glucose (Sigma–Aldrich) supplemented with 10% fetal calf serum (FCS). Culture supernatant was collected 2 days after transfection and passed through a 0.22-micron filter. The supernatant containing TMPRSS2 lentivector was added to human embryonic kidney cells, HEK293T, seeded on 6-well plates. Following 2 days of selection with 15 μg/ml of blasticidin (Invitrogen), the cells were incubated with the medium containing the ACE2 lentivector, and supplemented with 3 μg/ml puromycin (Sigma–Aldrich). After selection with puromycin, single cell lines were established by serial dilution. Both the double-transduced pool and the cloned cell lines were analyzed with SDS-PAGE using an anti-V5 antibody (#MA5-15253, Invitrogen) for TMPRSS2 and an anti-ACE2-antibody (#15983, Cell Signaling Technology).

HEK293T and HEK293T-ACE2-TMPRSS2 cells were maintained in DMEM supplemented with 10% FCS, 2% L-Glutamine, and 1% penicillin/streptomycin. Human lung epithelial cell line Calu-1was kept in Roswell Park Memorial Institute (RPMI) 1640 medium supplemented with 10% FCS, 1% L-Glutamine, and 1% penicillin/streptomycin. Another human lung epithelial cell line Calu-3 and human colorectal adenocarcinoma cell line Caco-2 were kept in Minimum Essential Medium Eagle (MEM) supplemented with 20% FCS, 1% L-Glutamine, 1% penicillin/streptomycin, and 1% MEM Non-essential Amino acid Solution (100X). VeroE6-TMPRSS2 cells (ATCC^®^ CRL-1586) were maintained in MEM supplemented with 10% FCS, 1% L-Glutamine and 1% penicillin/streptomycin. All the cell lines were incubated at 5% CO2 and 37°C.

We addressed the expression level of *ACE2, TMPRSS2, FURIN* and *GAPDH* with qPCR in all the cell lines (Fig. S1) by using human specific primers (Table S1).

### Production of luciferase-encoding lentiviral vector pseudotyped with WT and mutant SARS-CoV-2 S-glycoprotein, and VSV-glycoprotein

HEK293T cells grown in T175 flask were transfected by using TransIT-2020 reagent (Mirus Bio) with p8.9NDSB (Berthoux et al., 2004), pWPI-puro expressing *Renilla* luciferase, pEBB-GFP, and pCAGGS, an expression vector containing the SARS-CoV-2 spike glycoprotein cDNA of the Wuhan-Hu-1 reference strain (NC_045512.2). The last 18 codons of spike glycoprotein were deleted to enhance the plasma membrane transport. Pseudotyped viruses harbouring the glycoprotein of vesicular stomatitis virus (VSV-G) were produced following the same protocol by using VSV-G envelope expression vector pMD2.G (#12259, Addgene). The culture media was replaced 12-16 h after transfection with fresh DMEM High Glucose (Sigma– Aldrich) supplemented with 10% FBS. The supernatant containing the SARS-CoV-2 spike glycoprotein- or VSV-G-harboring pseudoviruses was collected 48 h after transfection, and passed through a 0.22-micron filter.

Mutations in RBD regions were generated by synthetic DNA (Integrated DNA Technologies) using PvuII and HpaI restriction sites. Lentiviral vectors pseudotyped with mutant spikes were produced by following a similar protocol as the wild type particles with minor changes. Instead of pWPI-puro expressing *Renilla* luciferase, pWPI plasmid carrying both GFP and *Renilla* luciferase were used to eliminate the use of pEBB-GFP plasmid.

### Detection of viral infection by pseudotyped SARS-CoV-2 or VSV-G viruses and native SARS-CoV-2

For the experiments with pseudotyped viruses, HEK293T-ACE2-TMPRSS2 cells were cultured in poly-L-1ysine coated 96-well plates (ViewPlate 96, PerkinElmer Life Sciences). Next day, the cells received varying concentrations of listed drugs (Table 1) and pseudotyped lentiviruses harboring S-protein of SARS-CoV-2 or glycoprotein of VSV. Following 24 h incubation with drugs and viral particles, cells were washed once with PBS. After cells were lysed for 15 min at RT, *Renilla* luciferase reporter was used to measure viral entry. For this purpose, luciferase activity was measured with a plate reader employing dispenser feature (Varioskan Flash, ThermoFisher Scientific) after substrate addition to each well *(Renilla* Luciferase Assay System, E2820, Promega or Coelenterazine native, cat#303, Nanolight Technology). In order to measure background signal, uninfected cells and empty wells were included into the assay plate.

All work with infectious SARS-CoV-2 virus was conducted in a Biosafety Level 3 (BSL-3) Laboratory of UH at Haartman Institute. SARS-CoV-2 virus isolates (wild type, B.1.1.7, and B.1.351), obtained from nasopharyngeal swabs of patients (Cantuti-Castelvetri et al., 2020; Virtanen et al., 2021) (MOI 0.05), were incubated with Calu-1 cells in 48-well plates (40.000 cells/well) for 1 h at 37°C and 5% CO2, after which virus inoculum was removed and cells were washed twice with PBS. The compounds were added to the cells in duplicates, diluted in virus growth medium (VGM) (RPMI-1640 supplemented with 2% FCS, L-glutamine, penicillin and streptomycin), 0.1% DMSO in VGM was used as control. Samples of the supernatant were collected at 1 h, 24 h and 48 h post infection for qRT-PCR and at 24 h and 48 h for the TCID50 assay. Viral RNA was extracted using RNeasy Mini Kit (Qiagen, Germany), and SARS-CoV-2 qRT-PCR was performed using primers, probe and an *in vitro* synthesized control for RNA-dependent RNA polymerase (RdRp) as described earlier (Corman et al., 2020; Lin et al., 2021). Infectious virus titers were determined by TCID50 measurement of VeroE6-TMPRSS2 cells (Virtanen et al., 2021). Shortly, 10-fold dilutions of the samples were inoculated to VeroE6-TMPRSS2 cells, incubated for 5 days, fixed with 10% formaldehyde for 30 min RT and stained with crystal violet.

A flow chart was prepared to summarize the timeline of experiments regarding the pseudotyped viruses and the native virus (Fig. 1).

**Figure 1.**
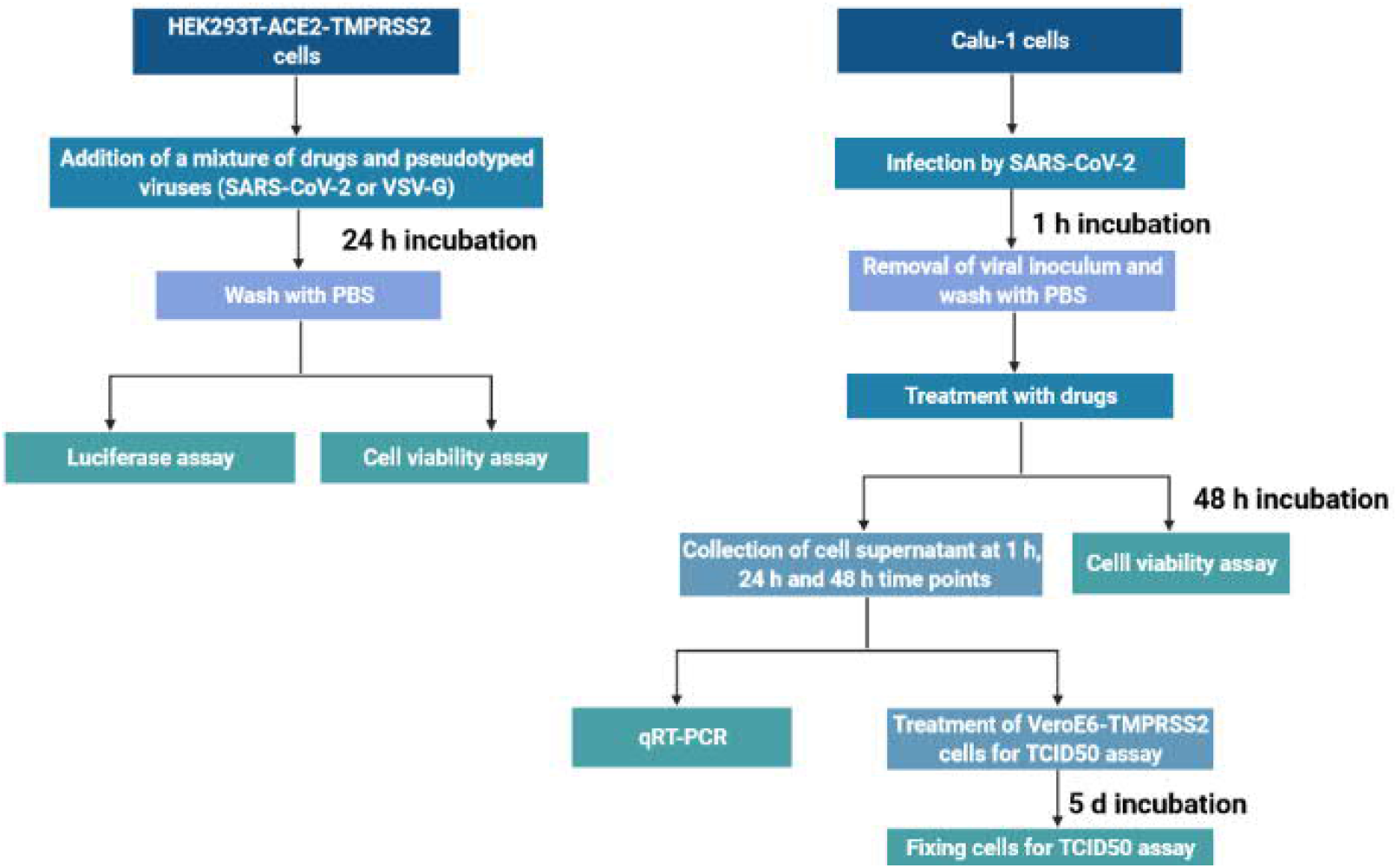
Protocols that were followed for infection, treatment, sample collection and assays.

### Cell viability assay

Cell culturing, treatment, and infection protocols were followed as described for HEK293T-ACE2-TMPRSS2 and Calu-1 cells (Fig. 1). At the end of 24 h or 48 h period, cells were washed once with PBS and their viability was measured with a kit designed to quantify ATP level according to instructions of the manufacturer (CellTiterGlo Luminescent Cell Viability Assay, Promega).

### Data and statistical analysis

#### Pseudotyped viruses

The number of samples per treatment group was determined based on our earlier studies (Casarotto et al., 2021; Fred et al., 2019). Each n represents an independent value. Data were normalized with control groups to avoid unwanted sources of variation, pooled together, and analyzed in GraphPad Prism 6.0 software. Pseudotyped virus experiments were analyzed by unpaired t-test or one-way analysis of variance (or their nonparametric equivalents) followed by Dunnett or Dunn’s *post hoc* tests, when appropriate. The luciferase activity readings from pseudotyped viruses (SARS-CoV-2 and VSV-G) were replotted together to run a comparative analysis. We compared the drug effect on the infection by SARS-CoV-2 spike and VSV-G pseudotyped viruses by two-way analysis of variance followed by Sidak’s multiple comparison test. The mutant pseudotyped virus experiments (infection) were analyzed by two-way analysis of variance followed by Sidak’s multiple comparison. IC50 values were calculated in GraphPad Prism software by using the non-linear curve fitting of the data with log(inhibitor) vs. response-Variable slope function after conversion of drug concentrations to logarithmic scale.

#### Native SARS-CoV-2 viruses

Due to limited resources for BSL-3 work with the native virus, experiments with SARS-CoV-2 was conducted only once with relatively smaller n. Experiments using native virus were analyzed by two-way analysis of variance followed by Dunnett or Sidak post hoc test in GraphPad Prism 6.0 or by multiple two-way analysis of variance test in JASP (0.14.1). The control group of some samples were the same, as they were tested in the same plate. Therefore, the same control group was represented in different plots. The plots representing changes in the genome copy number were plotted in linear scale, whereas TCID50 values were represented in log10 scale in GraphPad Prism.

For all the experiments, values of p< 0.05 were considered significant. Presence of any outliers was calculated by using Grubb’s test on GraphPad Prism website (https://www.graphpad.com/quickcalcs/grubbs1), and excluded from the analysis.

Statistical analysis were provided in Table S2 for the corresponding figures.

### Results

#### HEK293T-ACE2-TMPRSS2 cell line is responsive to camostat mesylate

Camostat mesylate, a clinically tested serine protease inhibitor, has been shown to inhibit the activity of TMPRSS2 and reduce the SARS-CoV-2 infection in cell lines expressing TMPRSS2 (Hoffmann, Hofmann-Winkler, Smith, et al., 2020; Hoffmann, Kleine-Weber, Schroeder, et al., 2020; Kawase et al., 2012). In order to confirm the activity of TMPRSS2 in the HEK293T-ACE2-TMPRSS2 cell line, we treated the cells with different doses of camostat mesylate for 24 h and measured the level of viral infection. The infection of HEK293T-ACE2-TMPRSS2 cell line by the pseudotyped lentiviruses was significantly reduced suggesting that that these cells express TMPRSS2 and the activity of this protease is blocked by camostat mesylate (Fig. S2).

#### Antidepressant drugs show antiviral activity against pseudotyped viruses and the first wave (B.1) lineage of SARS-CoV-2

HEK293T-ACE2-TMPRSS2 cells were treated with a mixture of pseudotyped viral particles and one of the following antidepressants: fluoxetine, citalopram, paroxetine, fluvoxamine, venlafaxine, reboxetine, clomipramine, imipramine, or desipramine (Table 1) for 24 h. We found that all the drugs significantly reduced the viral infection, as measured by luciferase reporter activity (Fig. 2). According to the cell viability assay where ATP level was measured in infected HEK293T-ACE2-TMPRSS2 cells, all the tested compounds, except venlafaxine (Fig. 2e) and reboxetine (Fig. 2f), were slightly toxic at higher concentrations (Fig. 2).

**Figure 2.**
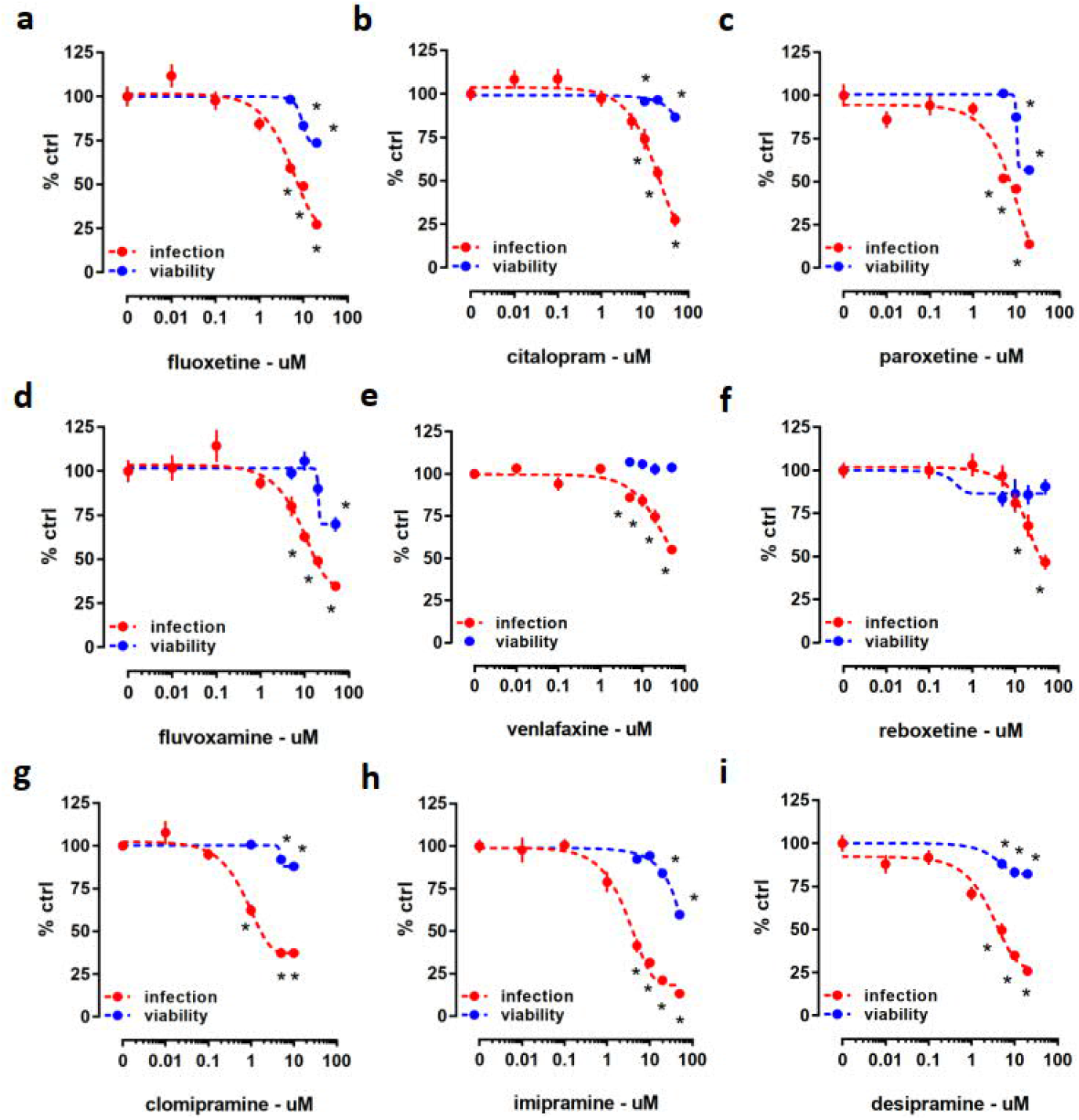
Luciferase reporter activity (infection) and ATP level (viability) in HEK293T-ACE2-TMPRSS2 cells challenged with SARS-CoV-2 pseudotyped viruses and pharmacologically diverse antidepressant drugs for 24 h. Treatment with (**a**) fluoxetine (IC50= 5.992 μM), (**b**) citalopram (IC50= 27.51μM), (**c**) paroxetine (IC50= 12.55 μM), (**d**) fluvoxamine (IC50= 10.54 μM), (**e**) venlafaxine (IC50= 36.35 μM), (**f**) reboxetine (IC50= 17.69 μM), (**g**) clomipramine (IC50= 0.75 μM), (**h**) imipramine (IC50= 3 μM), and (**i**) desipramine (IC50= 8.097μM) significantly reduced luciferase reporter activity. At higher concentrations, all the compounds, except (**e**) venlafaxine and (**f**) reboxetine reduced ATP levels in the luminescent cell viability assay after 24 h incubation. *p< 0.05 from control group (0). Data represented as mean ± SEM. SARS-CoV-2 infected Calu-1 cells were treated with fluoxetine, reboxetine, clomipramine, imipramine, citalopram or venlafaxine up to 48 h at concentrations of 5, 10, and 20 μM. The base level of viral RNA that was measured 1 h after drug treatment revealed no significant changes between control and treatment groups (Fig. 3). None of the compounds significantly changed viral replication or the amount of infectious virus after 24 h of treatment (Fig. 3). On the other hand, fluoxetine (Fig. 3a,b), reboxetine (Fig. 3c,d), clomipramine (Fig. 3e,f), and imipramine (Fig. 3g,h) were effective after 48 h to reduce the genome copies of SARS-CoV-2 and TCID50. However, citalopram (Fig. 3i,j) and venlafaxine (Fig. 3k,l) were ineffective on reducing viral replication and infectious particles of native SARS-CoV-2. Viability of Calu-1 cells was also measured in mock and infected cells after 48 h treatment with these compounds (Fig. S3). Fluoxetine and clomipramine reduced viability in Calu-1 cells (Fig. S3a,e). Other compounds, such as venlafaxine, did not have any significant effect on the viability of Calu-1 cells (Fig. S3c), although some differences were observed in mock vs. infected groups of citalopram and imipramine treated cells (Fig. S3b,f).

**Figure 3.**
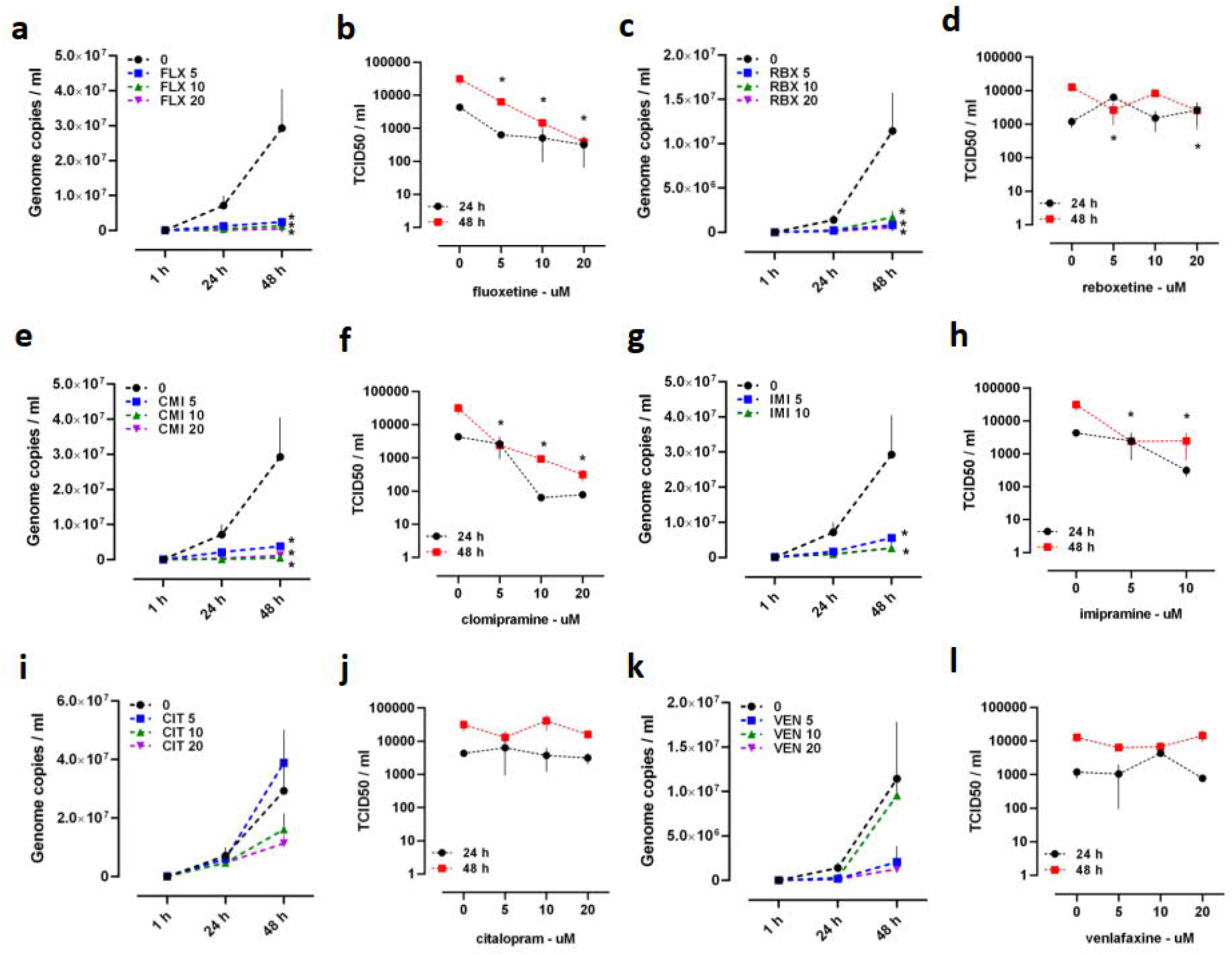
The effect of antidepressant drugs on the genome copies of SARS-CoV-2 (B.1) measured by qRT-PCR and on the amount of infectious virus detected by TCID50 assay in Calu-1 cells. (**a, b**) Fluoxetine, (**c, d**) reboxetine, (**e, f**) clomipramine, and (**g, h**) imipramine were ineffective after 24 h of treatment, but they reduced genome copy number of SARS-CoV-2 and the amount of infectious viral particles after 48 h of treatment. (**i, j**) Citalopram and (**k, l**) venlafaxine failed to significantly change the genome copy number of SARS-CoV-2 and TCID50. Genome copies were plotted in linear scale, while TCID50 values in log scale. Fluoxetine: FLX, Reboxetine: RBX, Clomipramine: CMI, Imipramine: IMI, Citalopram: CIT, Venlafaxine: VEN. *p< 0.05 from control group (0) at corresponding time point. Data represented as mean ± SEM.

#### Antipsychotics can decrease the viral infection

The antiviral activity of antipsychotics chlorpromazine, flupenthixol, and pimozide were tested in HEK293T-ACE2-TMPRSS2 cells against pseudotyped viruses harboring SARS-CoV-2 spike protein (Table 1). Following 24 h incubation, all of these compounds were able to prevent viral infection, although we also observed slight reduction of cell viability (Fig. 4).

**Figure 4.**
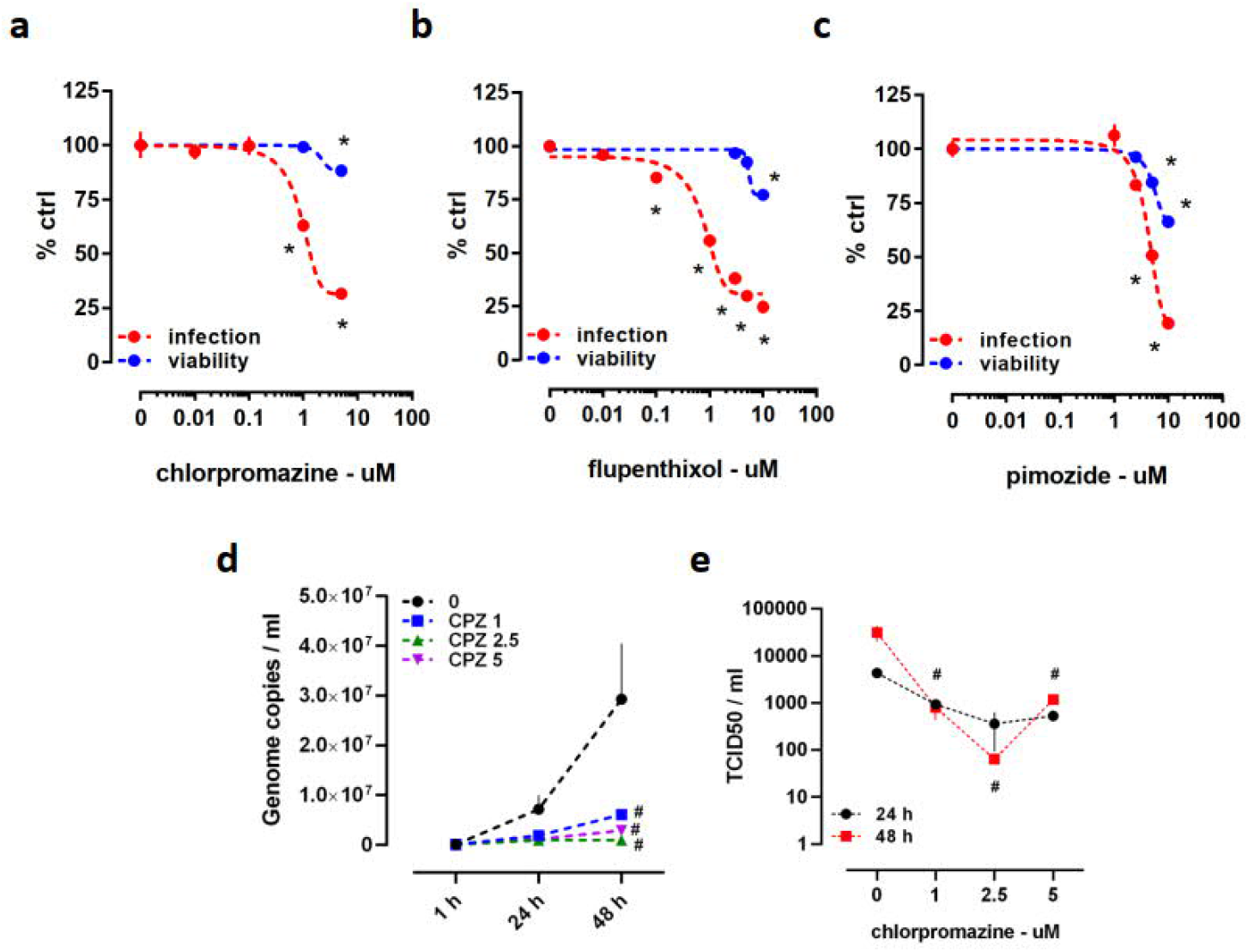
Luciferase reporter activity (infection) and ATP level (viability) in HEK293T-ACE2-TMPRSS2 cells challenged with SARS-CoV-2 pseudotyped viruses and antipsychotics. The effect of chlorpromazine against SARS-CoV-2 (B.1) in Calu-1 cells measured by qRT-PCR and TCID50 assay. Decline of luciferase reporter activity was observed at multiple doses of (**a**) chlorpromazine (IC50= 0.972 μM), (**b**) flupenthixol (IC50= 1.072 μM), and (**c**) pimozide (IC50= 4.539 μM) following 24 h incubation. Treatment of cells with (**a**) chlorpromazine, (**b**) flupenthixol, and (**c**) pimozide combined with viral infection decreased cell viability indicated by the ATP level. (**d**) Treatment with chlorpromazine for 48 h decreased the genome copy number of SARS-CoV-2 and (**e**) the amount of infectious virus. Genome copies were plotted in linear scale, while TCID50 values in log scale. Chlorpromazine: CPZ. #p< 0.05 from control group (0). *p< 0.05 from control group (0) at corresponding time point. Data represented as mean ± SEM.

Chlorpromazine (1, 2.5, and 5 μM) was also tested in SARS-CoV-2 (B.1 lineage) infected Calu-1 cells. Treatment for 48 h with all the tested concentrations of chlorpromazine were able to induce a significant reduction of total virus (Fig. 4d). In line with this, we observed diminished amount of infectious virus after 48 h treatment (Fig. 4e). No change in the viability of Calu-1 cells was observed in mock and infected cells after 48 h treatment with chlorpromazine (Fig. S3g).

### Some psychoactive drugs fail to prevent the infection by the pseudotyped viruses

We next tested whether ketamine and ketamine metabolites (2S,6S-HNK and 2R,6R-HNK) which have received attention during recent years because of their potential as rapid-acting antidepressant drugs (Abdallah et al., 2015; Zanos et al., 2018) might also inhibit infection by the pseudotyped virus. We found that neither ketamine nor the metabolites (Table 1) were effective reducing the viral infection within 24 h of incubation period (Fig. 5a-c). Furthermore, other classical antidepressant drugs, phosphodiesterase-4 inhibitor rolipram and monoamine oxidase inhibitor phenelzine (Table 1) also failed to prevent the viral entry (Fig. 5d,e). We also tested carbamazepine, an anticonvulsant, which is also used as mood stabilizer for the treatment of bipolar disorder (Mitchell & Malhi, 2002). This compound also failed to change the viral infection in ACE2/TMPRSS2-expressing HEK cells after 24 h of treatment (Fig. 4f).

**Figure 5.**
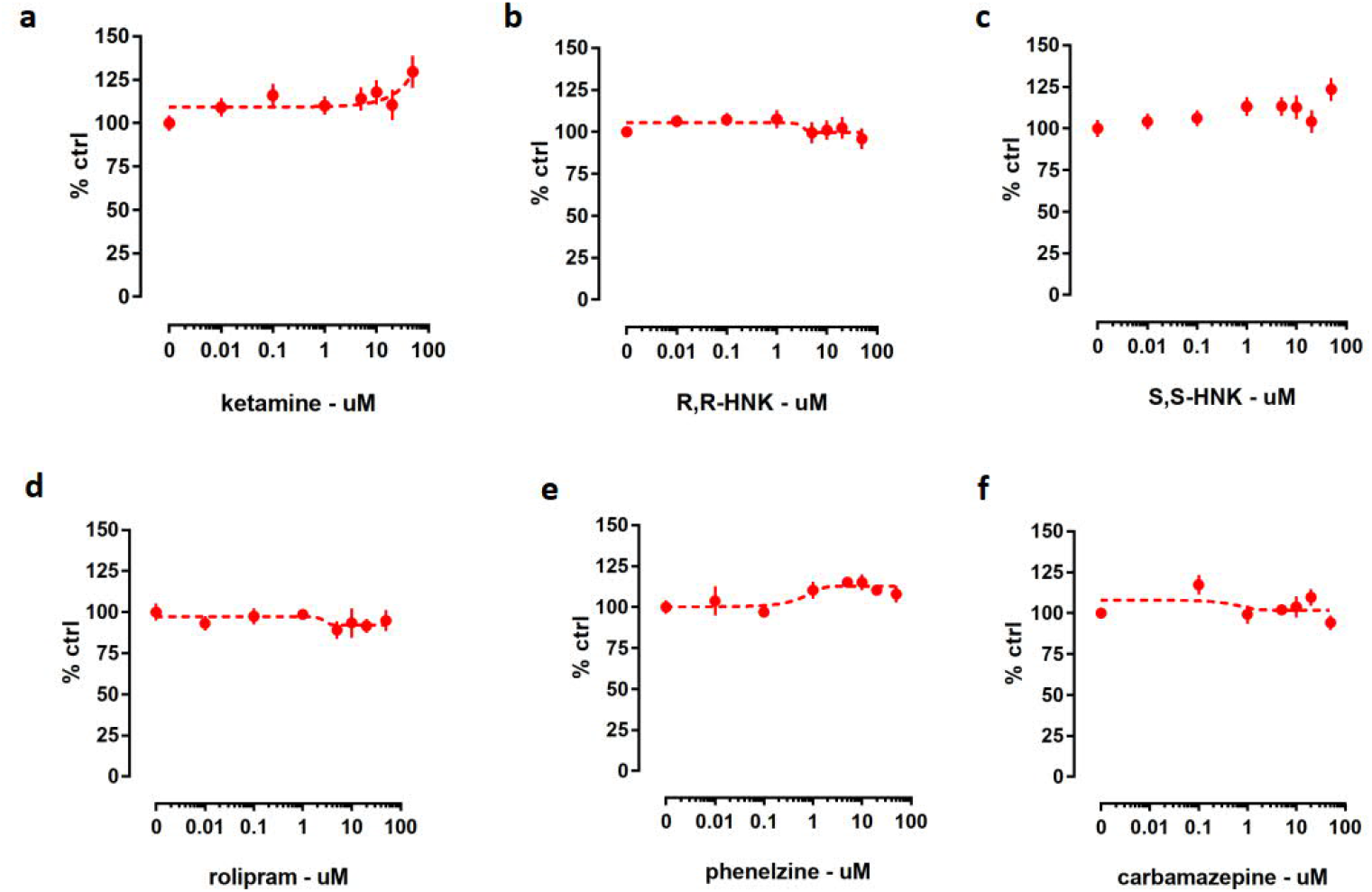
Luciferase reporter activity in HEK293T-ACE2-TMPRSS2 cells 24 h after infection by SARS-CoV-2 pseudotyped viruses and treatment with indicated compounds. (**a**) Ketamine and ketamine metabolites (**b**) 2R,6R-HNK and (**c**) 2S,6S-HNK did not change the luciferase reporter activity. Other classical antidepressant drugs (**d**) rolipram and (**e**) phenelzine were also found to be ineffective. (**f**) Anticonvulsant carbamazepine did not change the amount of luciferase activity. Data represented as mean ± SEM.

### Some of the tested drugs show SARS-CoV-2 spike specificity

We used pseudotyped viruses harboring glycoprotein of vesicular stomatitis virus (VSV) to address the specificity and potential viral-entry independent effects of tested compounds. HEK293T-ACE2-TMPRSS2 cells were treated with a mixture of VSV-G pseudotyped viruses and drugs including fluoxetine, citalopram, paroxetine, fluvoxamine, reboxetine, clomipramine, desipramine, chlorpromazine, flupenthixol, or pimozide for 24 h. Viral infection was quantified by measuring the luciferase activity (Figs. S4 and S5). While fluoxetine (Fig. S4a), clomipramine (Fig. S4c), chlorpromazine (Fig. S4e), flupenthixol (Fig. S4g), pimozide (Fig. S4i), and reboxetine (Fig. S4k) reduced the infection by the VSV-G pseudotyped viruses; citalopram (Fig. S5a), desipramine (Fig. S5c), paroxetine (Fig. S5e), and fluvoxamine (Fig. S5g) remained ineffective. Slight reduction of cell viability was observed in infected HEK293T-ACE2-TMPRSS2 cells after fluoxetine, clomipramine, chlorpromazine, pimozide, and reboxetine treatment (Fig. S4), while flupenthixol, citalopram, desipramine, paroxetine, or fluvoxamine did not exert a toxic effect (Fig. S4 and Fig. S5).

We compared the effectiveness of the tested antidepressants and antipsychotics against SARS-CoV-2 spike- and VSV-G-pseudotyped viruses. We compiled the data from these viruses, and found higher effectiveness of drugs in reducing the luciferase activity in SARS-CoV-2 pseudoviruses compared to VSV-G (Fig. S4b,d,f,h,j,l and S5b,d,f,h).

### Fluoxetine remains effective against pseudotyped viruses harboring the S protein RBD mutations and native variants VoC-1 (B.1.1.7) and VoC-2 (B.1.351)

Next, we addressed the effectiveness of fluoxetine against the pseudotyped viruses carrying mutations in the S protein that have been observed in some of the emerging SARS-CoV-2 variants (*Preliminary Genomic Characterisation of an Emergent SARS-CoV-2 Lineage in the UK Defined by a Novel Set of Spike Mutations - SARS-CoV-2 Coronavirus / NCoV-2019 Genomic Epidemiology*, 2020; Tegally et al., 2020; *Genomic Characterisation of an Emergent SARS-CoV-2 Lineage in Manaus*, 2021). We found that fluoxetine (10 μM) treatment for 24 h was effective against pseudotyped viruses carrying single point mutations in their S protein (N501Y, K417N, or E484K) (Fig. 6a, b), and also a triple mutant harboring combination of these three point mutations (N501Y/K417N/E484K) (Fig. 6c). Calu-1 cells which were infected with SARS-CoV-2 (B.1), VoC-1 (B.1.1.7) or VoC-2 (B.1.351) variants, and challenged with fluoxetine (10 μM) showed reduced amount of genome copies (Fig. 6d,e,f) and infectious virus (Fig. 6g,h,i). No difference was observed in the viability of Calu-1 cells infected with these variants and treated with fluoxetine for 48 h (Fig. S3h).

**Figure 6.**
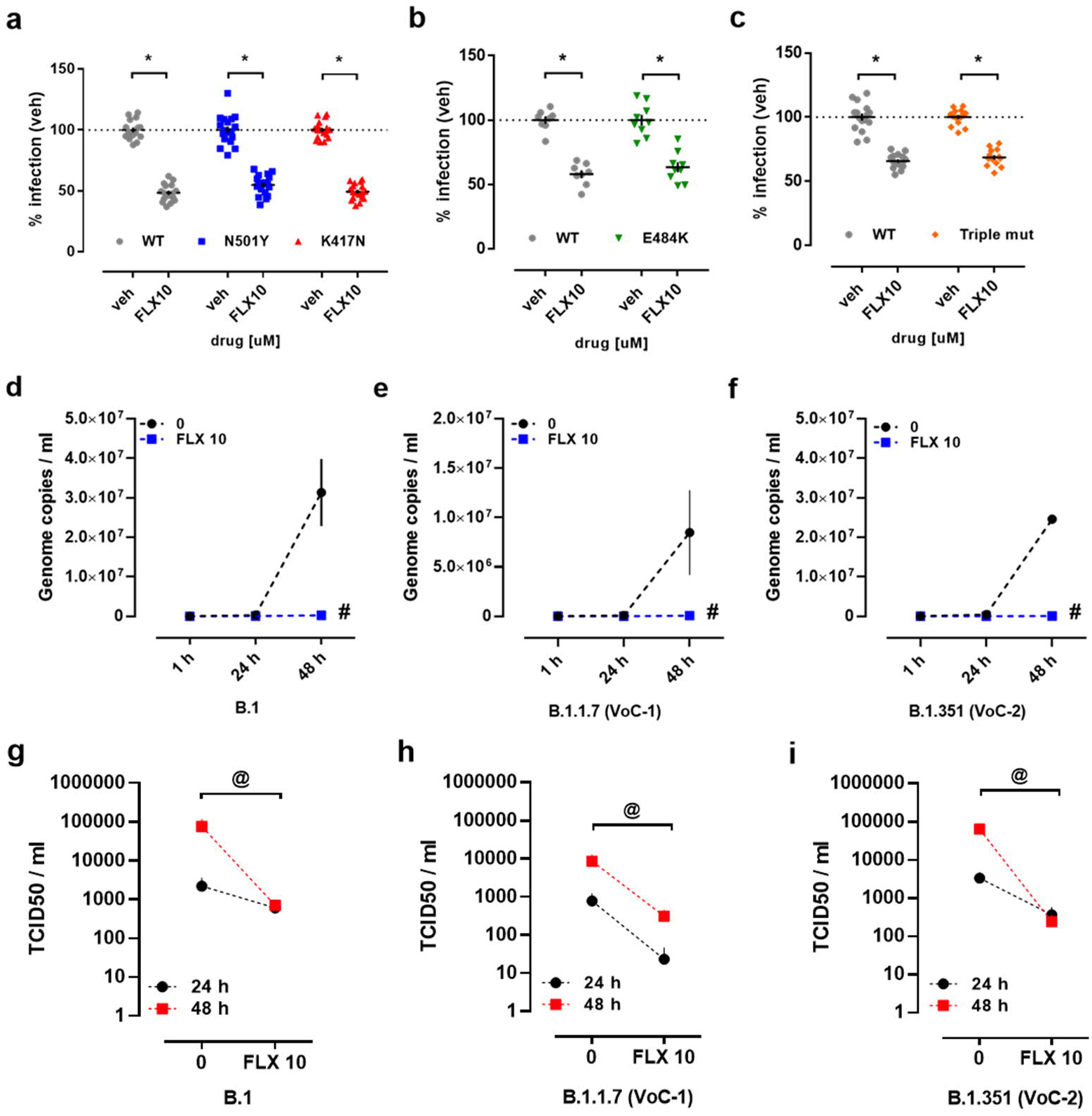
Luciferase reporter activity in HEK293T-ACE2-TMPRSS2 cells infected with mutant SARS-CoV-2 pseudotyped viruses and treated with fluoxetine. Antiviral activity of fluoxetine (10 μM) measured in Calu-1 cells infected with SARS-CoV-2 (B.1), B.1.1.7 (VoC-1), and B.1.351 (VoC-2) variants measured by qRT-PCR and TCID50 assay. Treatment with fluoxetine (10 μM, 24 h) significantly reduced the luciferase activity in cells infected with (**a**) WT (grey), N501Y (blue), K417N (red), (**b**) E484K (green) and (**c**) triple mutant (N501Y/K417N/E484K, orange), viral particles. Fluoxetine significantly reduced genome copy number and TCID50, respectively, in Calu-1 cells infected with (**d, g**) SARS-CoV-2 (B.1), (**e, h**) B.1.1.7 (VoC-1), and (**f, i**) B.1.351 (VoC-2). Fluoxetine: FLX, wild type: WT, variant of concern: VoC. #p<0.05 from control group (veh). *p<0.05 from control group (0). @p<0.05 from control group (0). Data represented as mean ± SEM.

## Discussion

In the present study, we identified the potential of some widely consumed antidepressant and antipsychotic drugs against the infection by SARS-CoV-2 (B.1) and the variants VoC-1 (B.1.1.7) and VoC-2 (B.1.351). The experiments using pseudotyped viruses to infect HEK293T cells overexpressing ACE2 and TMPRSS2 were aimed at identifying the direct effect of drugs on the entry phase of the viral infection, as we added the virus and drugs around the same time. The outcome measure by luciferase assay was a direct indication of the number of infected cells, due to replication-deficient nature of pseudotyped viruses. Drug concentrations required for the inhibition of SARS-CoV-2 infection are within the range of steady-state plasma concentrations assayed for these drugs (Bolo, 2000; Burke & Preskorn, 1999) indicating that the effects described here could be clinically meaningful.

By testing pseudotyped viruses harboring the VSV-G, we found that some compounds including fluoxetine, clomipramine, chlorpromazine, flupenthixol, pimozide, and reboxetine, counteract the infection in HEK-ACE-TMPRSS2 cells suggesting that these drugs are not specific for SARS-CoV-2 spike protein. In fact, it is not a surprising that these compounds are effective against the entry of other viruses, as this has been reported in studies testing clomipramine and flupenthixol against Ebola virus (Johansen et al., 2015). We observed that the tested compounds induce more dramatic effect on the entry of viruses pseudotyped with the spike protein of SARS-CoV-2 when compared to the ones harbouring the VSV glycoprotein. This speaks against the possibility that these drugs act at the level of luciferase transcription, as the impact they induce differs between pseudotyped viruses harbouring distinct viral entry proteins. Another interesting finding is that some drugs, such as, fluvoxamine, paroxetine, desipramine, and citalopram did not exert any effect on the infection by VSV-G particles suggesting the SARS-CoV-2 spike specificity of these compounds.

In experiments with human respiratory/lung epithelial cells and native virus, Calu-1 cells were first exposed to the SARS-CoV-2 virus, which was followed by removal of this inoculum and addition of drugs to be tested. In this scenario, drugs do not necessarily interfere with the initial infection of cells by the virus, which was at low multiplicity of infection (MOI 0.05) but with viral particle production and infection of the neighboring cells. The decrease in viral replication that we observed after treating the cells for 48 h with the tested drugs and the reduction of TCID50 as a secondary outcome could be caused by 1) the blockade of virus entry during the subsequent waves of Calu-1 cell infection; 2) a direct effect on the mechanisms of viral replication, packaging and release of new viral particles; 3) an indirect effect through the regulation of innate immunity; or 4) a combination of the above effects. The first alternative, that the drugs interfere with viral infection of Calu-1 cells later on when the cells release newly packed viruses to medium thereby explaining the decline of viral genome copies and TCID50, would also be compatible with our findings with the pseudotyped virus.

Recently, a number of studies presented evidence supporting the effects of antidepressant drugs and related psychoactive drugs as antiviral compounds against SARS-CoV-2 (Carpinteiro et al., 2020; Drayman et al., 2020; Hoertel et al., 2021; Lenze et al., 2020; M. Plaze et al., 2020; Marion Plaze et al., 2020; Schloer et al., 2020; Yang et al., 2020; Zimniak et al., 2021). One of these studies addressing SSRIs for their antiviral activity did not show any effect of escitalopram and paroxetine, while fluoxetine was successful with inducing antiviral activity after 3 days of treatment (Zimniak et al., 2021). Although this evidence supports our conclusion on the fluoxetine effect, paroxetine data remains controversial. We have also observed that citalopram significantly reduced the viral infection in HEK cells, while it remained ineffective against SARS-CoV-2 in Calu-1 cells. This piece of evidence suggests that HEK cells challenged with pseudotyped viruses and Calu-1 cells exposed to SARS-CoV-2 may recruit different machineries that the drugs can act on, which may explain this inconsistent data on the citalopram effect.

In another study, addressing the antiviral activity of fluoxetine, the same conclusion as ours suggested antiviral properties of this compound (Schloer et al., 2020). The authors have also described cholesterol-trapping and luminal pH altering effect of fluoxetine in endolysosomal compartments, which in the long term impairs the release of viral genetic material into the cytosol of target cells from endosomes (Schloer et al., 2020). The infection by SARS-CoV-2 activates acid sphingomyelinase (ASM) enzyme, leading to cell surface accumulation of ceramide, which in turn could facilitate the infection. However, the ASM activity can be inhibited by amitriptyline, imipramine, desipramine, fluoxetine, sertraline, escitalopram, and maprotiline, which reduced the infection by SARS-CoV-2 (Carpinteiro et al., 2020). Collectively, the data suggest that psychoactive drugs that have been categorized as functional inhibitors of acid sphingomyelinase activity (FIASMA) (Kornhuber et al., 2010, 2011) can significantly reduce the SARS-CoV-2 infection (Carpinteiro et al., 2020; Schloer et al., 2020). Most of the compounds we tested in the present study showed antiviral activity, and a number of them are considered FIASMAs. On the other hand, we have found that the non-FIASMA compounds venlafaxine and reboxetine (Kornhuber et al., 2011) also demonstrate antiviral activity. Moreover, citalopram, one of the compounds that was found to reduce the ASM activity, albeit at weaker level (Kornhuber et al., 2008), remains ineffective against native SARS-CoV-2 virus as shown by the present study and by others (Zimniak et al., 2021). Therefore, inhibition of ASM by these drugs can only partially explain the role of psychoactive drugs for the reduction of SARS-CoV-2 infection.

A recent study reported that the prevalence of COVID-19 was higher in health care professionals compared to patients in psychiatric ward of a hospital in Paris, which prompted the authors to suggest that the consumption of chlorpromazine protects against COVID-19 (M. Plaze et al., 2020; Marion Plaze et al., 2020). In a followup study, chlorpromazine was shown to exhibit antiviral activity against SARS-CoV-2 infection after 2 days of treatment of monkey and human cell lines which is in line with the clinical observation (Marion Plaze et al., 2020). Earlier studies on chlorpromazine have also demonstrated the antiviral activity of this compound against other coronaviruses (Cong et al., 2018; Dyall et al., 2014; Wilde et al., 2014). The action of chlorpromazine has been linked to inhibition of clathrin-mediated endocytosis which is possibly effective for the blockade of viral entry to target cells (Inoue et al., 2007; L. H. Wang et al., 1993; Weston et al., 2020). Although we did not address the mechanism of action, our data on the effect of chlorpromazine on HEK cells and Calu-1 cells are in agreement with these earlier studies. Other antipsychotics, flupenthixol and pimozide, identified as inhibitors of pseudotyped viral infection of HEK cells in our study, have also been confirmed by others as antiviral agents (Drayman et al., 2020; Yang et al., 2020). Pimozide, tested by computational docking analysis and *in vitro* assays, has been suggested to inhibit main protease of SARS-CoV-2 (M^Pro^) (Vatansever et al., 2020).

One of the challenges of drug repurposing is to find compounds that would achieve antiviral activity at safe concentrations for human use. Supporting our *in vitro* findings, Hoertel and colleagues administered different classes of antidepressants to 460 COVID-19 patients after admission to hospital. Administration of these drugs, particularly fluoxetine, escitalopram, paroxetine, and venlafaxine at an average of 20 days with fluoxetine-equivalent dose of 21.6 mg was associated with decreased risk of intubation or death by COVID-19 (Hoertel et al., 2021). Moreover, treatment of outpatients with a positive COVID-19 test with the SSRI fluvoxamine for 2 weeks was effective in reducing the development of clinical deterioration compared to the placebo group (Lenze et al., 2020). Despite not addressing it directly, the authors suggest that the antiviral activity of fluvoxamine might be due to the Sigma-1 receptor (S1R) agonism by this compound, as the activation of this receptor is known to regulate cytokine production (Lenze et al., 2020). Not only fluvoxamine, but also many other psychoactive drugs bind to S1R, including sertraline, fluoxetine, citalopram, imipramine, paroxetine, and desipramine (Narita et al., 1996). However, venlafaxine, one of the compounds that showed antiviral activity in our study in HEK cells, has a very weak affinity to this chaperone (Ishima et al., 2014). Moreover, ketamine can bind to both S1R and S2R (Robson et al., 2012), but this drug was ineffective as an antiviral reagent in the present study. Therefore, S1R agonism and the regulation of cytokine levels through this chaperon does not appear to fully explain how these drugs act as antiviral reagents.

One aspect that requires special attention is the cell type used in drug screening assays. Hoffmann and colleagues have shown that one of the most debated drugs recently, chloroquine, fails to inhibit SARS-CoV-2 infection of Vero cells expressing TMPRSS2 while lack of TMPRSS2 makes this drug effective as antiviral reagent (Hoffmann, Mösbauer, Hofmann-Winkler, et al., 2020). This finding suggests that the presence of certain proteins associated with viral entry, such as TMPRSS2, can render a drug effective or ineffective, and are required to maintain wild type nature of the infecting viral strains (Cantuti-Castelvetri et al., 2020). Therefore, it is important to implement cell lines that are phenotypically close to human respiratory epithelial cells to drug repurposing studies and validate the findings by different methodologies in order to increase the translational value (Hoffmann, Mösbauer, Hofmann-Winkler, et al., 2020). In the present study, we manipulated regular HEK293T cells to stably express ACE2 and TMPRSS2, initially identified targets that are important for SARS-CoV-2 infection (Hoffmann, Kleine-Weber, Schroeder, et al., 2020), and infection with the native SARS-CoV-2 virus was tested in a relevant human lung cell line, Calu-1, infected with wild-type low-passage viral strains (Cantuti-Castelvetri et al., 2020).

Emergence of the novel variants have raised a concern on the effectiveness of currently available SARS-CoV-2 vaccines for the neutralization of these variants (Callaway, 2021; Callaway & Ledford, 2021; Fontanet et al., 2021). Recent *in vitro* studies reported that some of these vaccines remain effective against the variants carrying a set of mutations (Muik et al., 2021; K. Wu et al., 2021; Xie et al., 2021); however, others have shown decreased effectiveness against these variants, particularly to those carrying the E484K mutation (Callaway & Mallapaty, 2021; Collier et al., 2021; Wadman, 2020; P. Wang et al., 2021). Our present data suggest that fluoxetine remains effective against the pseudotyped viruses harboring single mutations (N501Y, K417N, E484K), or a triple mutation (N501Y/K417N/E484K) as present in VoCs 2 and 3. In line with this, inhibitory effect of fluoxetine persists against the SARS-CoV-2 variants VoC-1 (B.1.1.7) and VoC-2 (B.1.351). Therefore, it is plausible to argue that antidepressants, as well as related-psychoactive compounds can be considered as an alternative treatment method for people infected with these variants of SARS-CoV-2.

Since the emergence of COVID-19, substantial progress has been made towards understanding the life cycle of SARS-CoV-2 and particularly how this virus can enter target cells. Dissecting the viral entry routes provides the opportunity to come up with alternative strategies for prevention and management of the viral infection. Studies that are currently available on the antiviral action of antidepressant drugs focus on the effect of these drugs at late entry (in cytoplasm of host cell) (Carpinteiro et al., 2020; Schloer et al., 2020; T. Tang et al., 2020). However, the effect of these drugs at the early entry pathway level (at host plasma membrane level) should also be addressed. Thermal shift assays followed by X-ray diffraction studies suggested specific sites of interaction for drugs, such as imipramine, clomipramine, thioridazine, paroxetine, and sertraline, on EBOV glycoprotein (Ren et al., 2018; Y. Zhao et al., 2018). Similar strategies can be followed to address whether these drugs can bind to SARS-CoV-2 S protein directly. Altogether, our data together with other recent studies suggest that safe and clinically available antidepressants drugs might become an additional tool for our fight against the COVID-19 pandemia.

## Acknowledgements

We thank Seija Lågas, Sulo Kolehmainen and Mira Utriainen for technical assistance. We extend our gratitude to Leonora Szirovicza for maintaining Calu-3 and Caco-2 cell lines. Dr. Emmy Verschuren is thanked for providing the Calu-1 cell line, Dr. Mirja Puolakkainen for Calu-3 cell line, Prof. Carl-Henrik von Bonsdorf for Caco-2 cell line, and Dr. Didier Trono for plasmids that were acquired through Addgene.

## Author Contributions

SMF, PCC and EC conceptualization; SMF, SK, HU, LL investigation; SMF, SK, KS, HU writing first draft, reviewed by SMF, SK, KS, OV, PCC, and EC.

