## supplement for "Antidepressant and antipsychotic drugs reduce viral infection by SARS-CoV-2 and fluoxetine show antiviral activity against the novel variants *in vitro*"

### Supplement Material

#### mRNA isolation, cDNA synthesis and quantitative PCR (qPCR)

Cells were washed with PBS (1X) once and collected after incubation in Qiazol lysis reagent (QIAGEN). Lysates were incubated in chloroform at RT for 3 min and vortexed rigorously for 15 seconds. After centrifugation at 12000 rpm for 10 min, aqueous phase was transferred to clean tubes and mixed with isopropanol. The samples were centrifuged for 10 min and the supernatant was discarded. Extensive washes in 70% ethanol was followed by a final wash in 96% ethanol. Pellets were air-dried and dissolved in RNAse-free MQ. The quantity and purity of RNA samples were measured with Nanodrop (Thermo Scientific).

Maxima First Strand cDNA Synthesis Kit for RT-qPCR with dsDNAse (#K1672, Thermo Scientific) was used to synthesize cDNA according to the instructions of the manufacturer. Genomic DNA contamination was assessed by including an "RT-" control and the reagent contamination was addressed by using a "no template" control.

Human-specific primers for qPCR were retrieved from the literature (Table S1). Maxima®SYBR Green qPCR Master Mix (2X) (Thermo Scientific, #K0253) was mixed with cDNA samples according to the instructions of the manufacturer and loaded to Hard-Shell® 96-well PCR plate (BioRad) as duplicates. Thermal cycler (BioRad CFX96 Real-Time System) was used for the assay with 44 cycles of amplification. Amplicon size and quality were verified by using agarose gel electrophoresis (data not shown).

**Table S1.** Human primers used for qPCR

| Primer Sequence (5' > 3') | Gene name | Amplicon Size (bp) | Reference |
|---|---|---|---|
| F: GGGATCAGAGATCGGAAGAAGAAA<br>R: AGGAGGTCTGAACATCATCAGTG | *ACE2* (human) pair 1 | 124 | (Ma et al., 2020) |
| F: AAACATACTGTGACCCCGCAT<br>R: CCAAGCCTCAGCATATTGAACA | *ACE2* (human) pair 2 | 199 | (Ma et al., 2020) |
| F: AATCGGTGTGTTCGCCTCTAC<br>R: CGTAGTTCTCGTTCCAGTCGT | *TMPRSS2* (human) pair 1 | 106 | (Ma et al., 2020) |
| F: CACTGTGCATCACCTTGACC<br>R: ACACACCGATTCTCGTCCTC | *TMPRSS2* (human) pair 2 | 196 | (Esumi et al., 2015) |
| F: GAGTCAACGGATTTGGTCGT<br>R: GACAAGCTTCCCGTTCTCAG | *GAPDH* (human) | 185 | (Yin et al., 2020) |
| F: CCTTCTTCCGTGGGGTTAG<br>R: GCAGTTGCAGCTGTCATGTT | *FURIN* (human) | 98 | (Yin et al., 2020) |

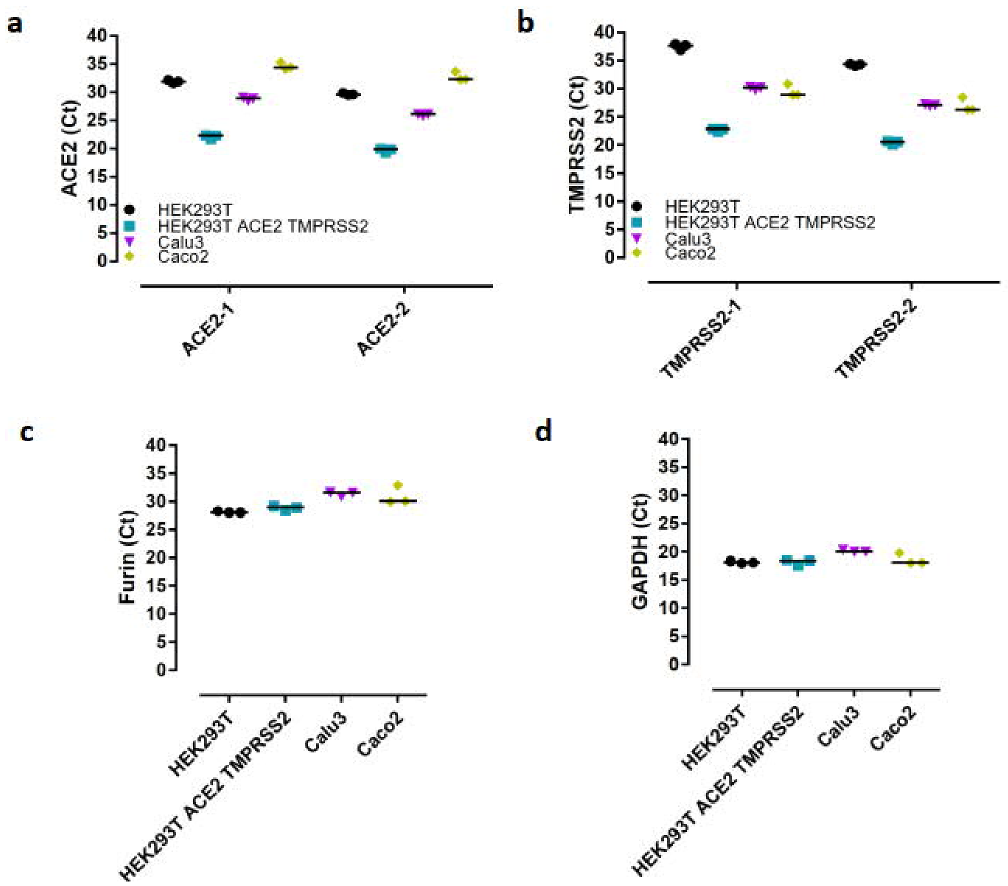


**Figure S1.** qPCR validation of *ACE2, TMPRSS2, FURIN* and the housekeeping gene *GAPDH* expression in cell lines. cDNA samples collected from HEK293T, HEK293T-ACE2-TMPRSS2, Calu-3, and Caco-2 cell lines were subjected to qPCR in order to address the expression of genes which are essential for the binding, processing, and entry of SARS-CoV-2. Ct values were represented for **(a)** ACE2 by using two sets of primers, **(b)** TMPRSS2 by using two sets of primers, **(c)** FURIN, and **(d)** GAPDH. n= 3 for all the groups.

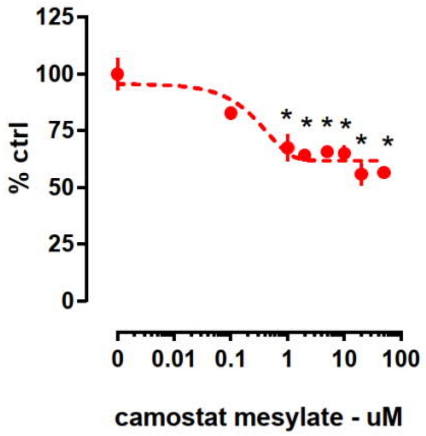


**Figure S2. Luciferase reporter activity in HEK293T-ACE2-TMPRSS2 cells challenged with pseudotyped viruses harboring SARS-CoV-2 spike protein and camostat mesylate for 24 h.** Camostat mesylate induces a reduction of luciferase activity in HEK 293T-ACE2-TMPRSS2 cells. *, $p < 0.05$ for comparison with the control group (0). Data represented as mean ± S.E.M.

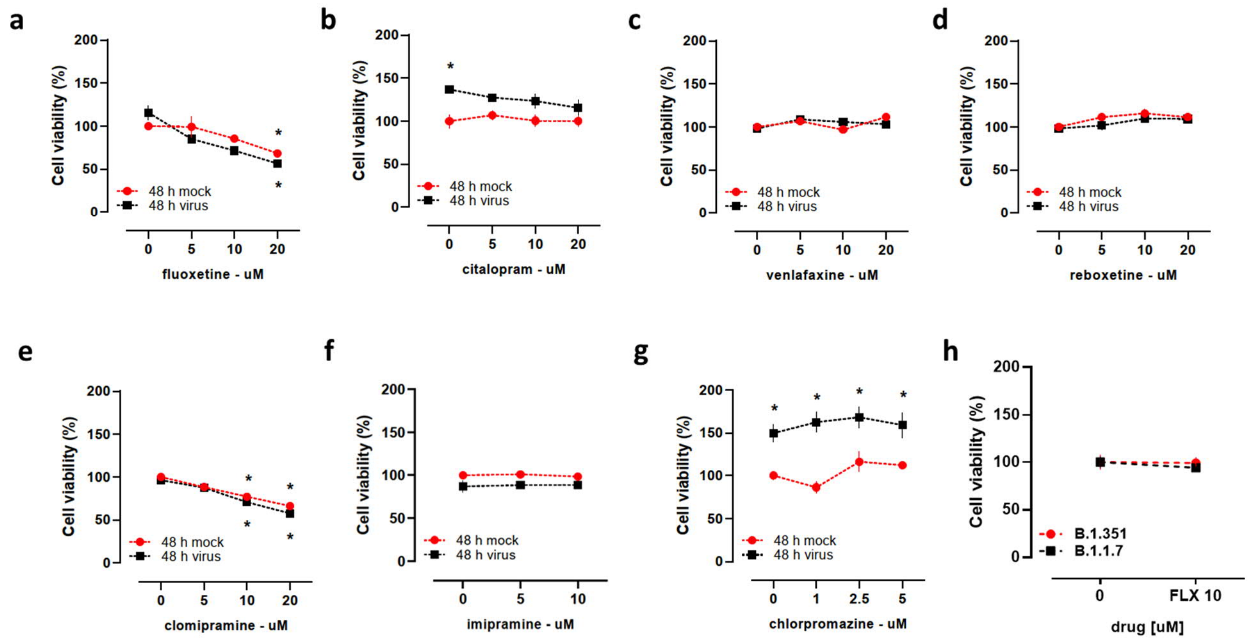


**Figure S3. Assessment of cell viability in non-infected and infected Calu-1 cells combined with 48 h drug treatment. (a)** Fluoxetine reduced the viability of Calu-1. **(b)** Citalopram, did not interfere with cell viability, while a slight increase was observed in the infected control group. **(c)** Venlafaxine did not induce any effect. **(d)** Reboxetine showed a slight potentiating effect. **(e)** Clomipramine decreased the cell viability in both non-infected and infected Calu-1 cells. **(f)** Imipramine itself did not affect the viability, while the viral infection induced slight reduction of viability in this batch of cells. **(g)** Chlorpromazine did not change the viability of Calu-1 cells, while viral infection increased the cell viability in this batch of cells. **(h)** Viability of cells infected by B.1.1.7 and B.1.351 variants of SARS-CoV-2 were not affected by fluoxetine (10 µM) treatment. *, $p < 0.05$ for comparison with the control group (48 h mock - 0). Data represented as mean ± S.E.M.

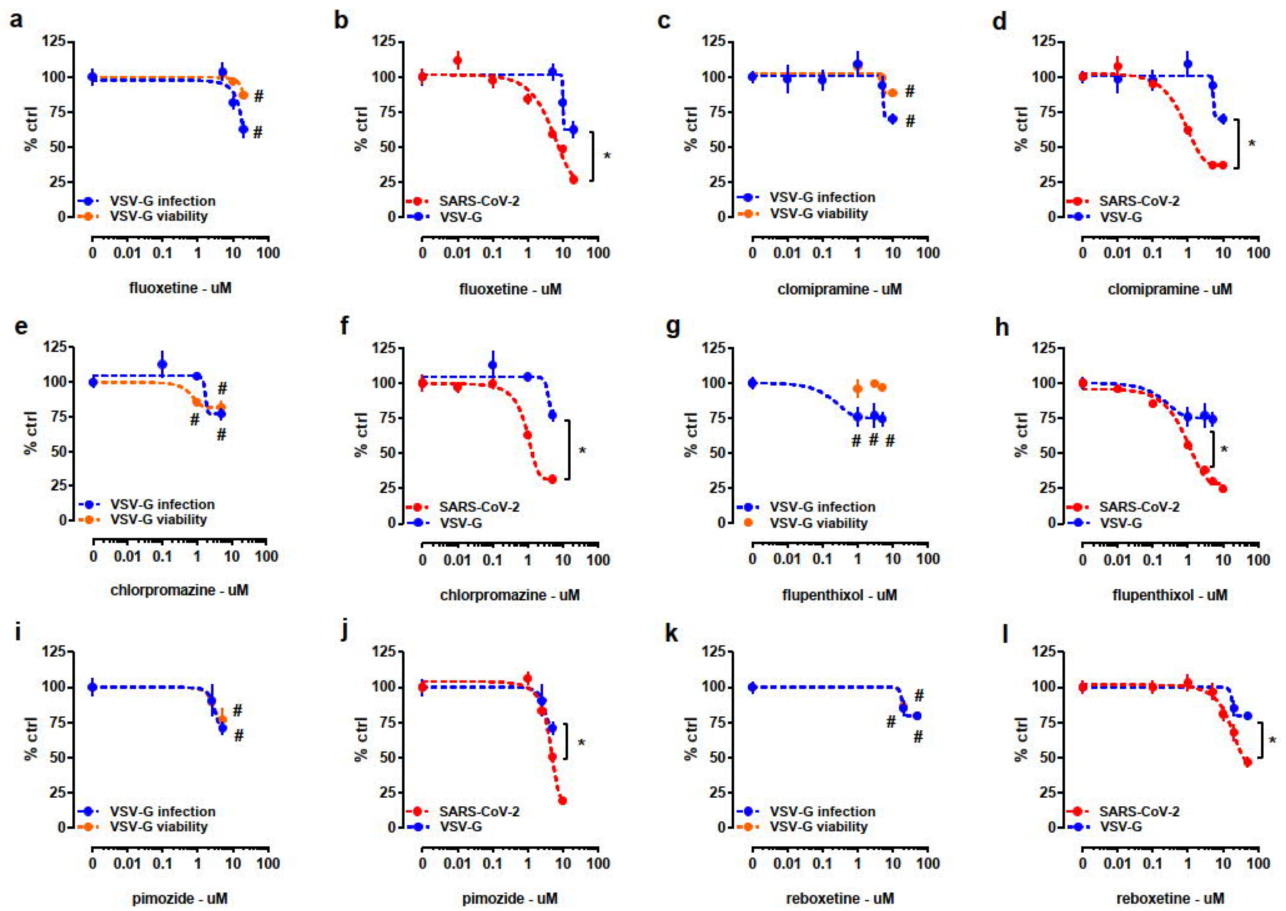


**Figure S4. Luciferase reporter activity and ATP level in HEK293T-ACE2-TMPRSS2 cells challenged with VSV-G pseudotyped viruses and antidepressant-antipsychotic drugs. Comparison of luciferase reporter activity in HEK cells infected with SARS-CoV-2 spike- and VSV-G-pseudotyped viruses.** 24 h treatment with **(a)** fluoxetine, **(c)** clomipramine, **(e)** chlorpromazine, **(g)** flupenthixol, **(i)** pimozide, and **(k)** reboxetine reduced the luciferase reporter activity (blue dashed line). Decrease of ATP level was observed following 24 h treatment with **(a)** fluoxetine, **(c)** clomipramine, **(e)** chlorpromazine, **(i)** pimozide, and **(k)** reboxetine indicating the reduction of cell viability in infected cells (orange dashed line or full circles). **(g)** Flupenthixol did not alter the viability of cells (orange full circles). Treatment with **(b)** fluoxetine (5, 10, 20 µM), **(d)** clomipramine (1, 5, 10 µM), **(f)** chlorpromazine (1 and 5 µM), **(h)** flupenthixol (1, 3, 5 µM), **(j)** pimozide (5 µM), and **(l)** reboxetine (20 and 50 µM) were more effective against the infection by SARS-CoV-2 spike pseudotyped viruses (red dashed line) than VSV-G pseudotyped viruses (blue dashed line). #$p < 0.05$ from control group (0), *$p < 0.05$ SARS-CoV-2 vs. VSV-G. Data represented as mean ± SEM.

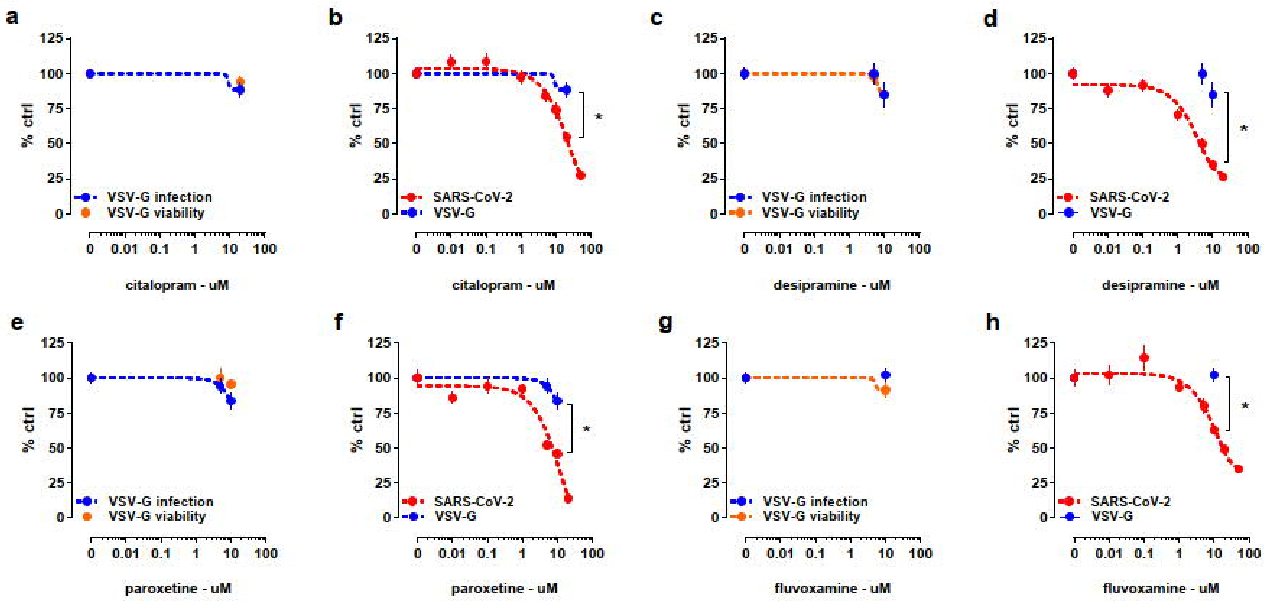


**Figure S5. Luciferase reporter activity and ATP level in HEK293T-ACE2-TMPRSS2 cells challenged with VSV-G pseudotyped viruses and antidepressant-antipsychotic drugs. Comparison of luciferase reporter activity in HEK cells infected with SARS-CoV-2 spike- and VSV-G-pseudotyped viruses.** 24 h treatment with **(a)** citalopram, **(c)** desipramine, **(e)** paroxetine, and **(g)** fluvoxetine did not significantly alter the luciferase reporter activity (blue dashed line or full circles). No significant change was observed in ATP level after treatment with **(a)** citalopram, **(c)** desipramine, **(e)** paroxetine, and **(g)** fluvoxamine indicating unaltered cell viability in infected cells (orange dashed line or full circles). Treatment with **(b)** citalopram (20 µM), **(d)** desipramine (5 and 10 µM), **(f)** paroxetine (5 and 10 µM), and **(h)** fluvoxamine (10 µM) were more effective against the infection by SARS-CoV-2 spike pseudotyped viruses (red dashed line) than VSV-G pseudotyped viruses (blue dashed line or full circle). *p< 0.05 SARS-CoV-2 vs. VSV-G. Data represented as mean ± SEM.

**Table S2.** Statistical analysis of the data.

| Graph | Test | N per group |
|---|---|---|
| 2a | Fluoxetine<br>One-way ANOVA:<br>Luciferase assay: Kruskal-Wallis (6) = 107.3, p< 0.0001*<br>Cell viability: F(3,20)= 134.6, p< 0.0001* | Luciferase assay: n= 26, 19, 18, 20, 20, 20, 24<br>Cell viability: n= 6/group |
| 2b | Citalopram<br>One-way ANOVA:<br>Luciferase assay: F(7,148)= 30.84, p< 0.0001*<br>Cell viability: F (3, 20)= 23.11, p< 0.0001* | Luciferase assay: n= 22, 16, 16, 22, 22, 22, 18, 18<br>Cell viability: n= 6/group |
| 2c | Paroxetine<br>One-way ANOVA:<br>Luciferase assay: Kruskal-Wallis (6) = 65.37, p< 0.0001*<br>Cell viability: F(3,16)= 98.75, p< 0.0001* | Luciferase assay: n= 12/group<br>Cell viability: n= 5/group |
| 2d | Fluvoxamine<br>One-way ANOVA:<br>Luciferase assay: Kruskal-Wallis (7) = 91.16, p< 0.0001*<br>Cell viability: F(4,25)= 8.047, p= 0.0003* | Luciferase assay: n= 24, 14, 14, 14, 14, 14, 18, 20<br>Cell viability: n= 6/group |
| 2e | Venlafaxine<br>One-way ANOVA:<br>Luciferase assay: F(7,87)= 24.08, p< 0.0001*<br>Cell viability: F(4,20)= 1.058, p= 0.4030 | Luciferase assay: n=12/group<br>Cell viability: n= 5/group |
| 2f | Reboxetine<br>One-way ANOVA:<br>Luciferase assay: F(6,77)= 14.30, p< 0.0001*<br>Cell viability: F(4,25)= 1.377, p= 0.2702 | Luciferase assay: n= 12/group<br>Cell viability: n= 6/group |
| 2g | Clomipramine<br>One-way ANOVA:<br>Luciferase assay: Kruskal-Wallis (5) = 81.31, p< 0.0001*<br>Cell viability: F(3,20)= 16.16, p< 0.0001* | Luciferase assay: n= 24, 12, 12, 18, 18, 17<br>Cell viability: n= 6/group |
| 2h | Imipramine<br>One-way ANOVA:<br>Luciferase assay: Kruskal-Wallis (7) = 75.25, p< 0.0001*<br>Cell viability: Kruskal-Wallis (4)= 26.58, p< 0.0001* | Luciferase assay: n= 18, 6, 6, 12, 12, 12, 12, 12<br>Cell viability: n= 6/group |
| 2i | Desipramine<br>One-way ANOVA:<br>Luciferase assay: Kruskal-Wallis (6) = 66.85, p< 0.0001* | Luciferase assay: n= 12/group<br>Cell viability: n= 5/group |

| | | |
|---|---|---|
| | Cell viability: F(3,16)= 16.48, p< 0.0001* | |
| 3a | Fluoxetine<br>Two-way ANOVA:<br>Time: F(2,12)= 6.669, p= 0.0113*<br>Treatment: F(3,12)= 8.885, p= 0.0022*<br>Interaction: F(6,12)= 4.779, p= 0.0103* | n= 2/group |
| 3b | Fluoxetine<br>Two-way ANOVA:<br>Time: F(1,8)= 8.614, p= 0.0189*<br>Treatment: F(3,8)= 8.181, p= 0.0081*<br>Interaction: F(3,8)= 4.852, p= 0.0329* | n = 2/group |
| 3c | Reboxetine<br>Two-way ANOVA:<br>Time: F(2,23) = 8.742, p= 0.0015*<br>Treatment: F(3,23)= 6.552, p= 0.0023*<br>Interaction: F(6,23)= 4.717, p= 0.0029* | 1 h: 0 (n= 3), 5 (n= 3), 10 (n= 2), 20 (n= 3)<br>24 h: 0 (n= 3), 5 (n= 3), 10 (n= 3), 20 (n= 3)<br>48 h: 0 (n= 3), 5 (n= 3), 10 (n= 3), 20 (n= 3) |
| 3d | Reboxetine<br>Two-way ANOVA:<br>Time: F(1,13)= 7.267, p= 0.0183*<br>Treatment: F(3,13)= 1.901, p= 0.1794<br>Interaction: F(3,13)= 6.307, p= 0.0071* | 24 h: 0 (n= 3), 5 (n= 2), 10 (n= 3), 20 (n= 2)<br>48 h: 0 (n= 3), 5 (n= 2), 10 (n= 3), 20 (n= 3) |
| 3e | Clomipramine<br>Two-way ANOVA:<br>Time: F(2,12)= 7.144, p= 0.0090*<br>Treatment: F(3,12)= 8,629, p= 0.0025*<br>Interaction: F(6,12)= 4.683, p= 0.0112* | n= 2/group |
| 3f | Clomipramine<br>Two-way ANOVA:<br>Time: F(1,7)= 4.477, p= 0.0721<br>Treatment: F(3,7)= 7.078, p= 0.0159*<br>Interaction: F(3,7)= 4.519, p= 0.0459* | 24 h: 0 (n= 2), 5 (n= 2), 10 (n= 2), 20 (n= 2)<br>48 h: 0 (n= 2), 5 (n= 2), 10 (n= 1), 20 (n= 2) |
| 3g | Imipramine<br>Two-way ANOVA:<br>Time: F(2,9)= 8.164, p= 0.0095*<br>Treatment: F(2,9)= 7.005, p= 0.0146*<br>Interaction: F(4,9)= 3.802, p= 0.0446* | n= 2/group |
| 3h | Imipramine<br>Two-way ANOVA:<br>Time: F(1,6)= 6.086, p= 0.0487*<br>Treatment: F(2,6)= 7.296, p= 0.0247*<br>Interaction: F(2,6)= 4.893, p= 0.0549 | n= 2/group |
| 3i | Citalopram<br>Two-way ANOVA:<br>Time: F(2,12)= 23.30, p< 0.0001*<br>Treatment: F(3,12)= 2.243, p= 0.1358<br>Interaction: F(6,12)= 1.838, p= 0.1739 | n= 2/group |
| 3j | Citalopram<br>Two-way ANOVA:<br>Time: F(1,8) = 10.21, p= 0.0127*<br>Treatment: F(3,8)= 0.9518, p= 0.4605<br>Interaction: F (3, 8) = 1.127, p = 0.3945 | n= 2/group |
| 3k | Venlafaxine<br>Two-way ANOVA:<br>Time: F(2,23)= 5.409, p= 0.0119*<br>Treatment: F(3,23)= 1.330, p= 0.2890<br>Interaction: F (6,23)= 1.000, p= 0.4488 | 1 h: 0 (n= 3), 5 (n= 3), 10 (n= 3), 20 (n= 3)<br>24 h: 0 (n= 3), 5 (n= 3), 10 (n= 3), 20 (n= 3)<br>48 h: 0 (n= 3), 5 (n= 2), 10 (n= 3), 20 (n= 3) |
| 3l | Venlafaxine<br>Two-way ANOVA:<br>Time: F(1,12)= 22.95, p= 0.0004*<br>Treatment: F(3,12)= 0.9331, p= 0.4547<br>Interaction: F(3,12)= 2.475, p= 0.1114 | 24 h: 0 (n= 3), 5 (n= 2), 10 (n= 3), 20 (n= 2)<br>48 h: 0 (n= 3), 5 (n= 2), 10 (n= 3), 20 (n= 2) |
| 4a | Chlorpromazine<br>One-way ANOVA:<br>Luciferase assay: Kruskal-Wallis (4) = 45.21, p< 0.0001*<br>Cell viability: F(2,21)= 42.42, p< 0.0001* | Luciferase assay: n= 20, 8, 8, 14, 14<br>Cell viability: n= 8/group |
| 4b | Flupenthixol<br>One-way ANOVA:<br>Luciferase assay: F(6,77)= 149.3, p< 0.0001*<br>Cell viability: F(3,16)= 19.04, p< 0.0001* | Luciferase assay: n= 12/group<br>Cell viability: n= 5/group |
| 4c | Pimozide<br>One-way ANOVA:<br>Luciferase assay: Kruskal-Wallis (4) = 50.77, p< 0.0001*<br>Cell viability: F(3,16)= 77.59, p< 0.0001* | Luciferase assay: n= 12/group<br>Cell viability: n= 5/group |
| 4d | Chlorpromazine<br>Two-way ANOVA: | n= 2/group |

| | | |
|---|---|---|
| | Time: F(2,12)= 8.987, p= 0.0041*<br>Treatment: F(3,12)= 7.656, p= 0.0040*<br>Interaction: F(6,12)= 4.226, p= 0.0162* | |
| 4e | Chlorpromazine<br>Two-way ANOVA:<br>Time: F(1,6)= 3.388, p= 0.1153<br>Treatment: F(3,6)= 6.248, p= 0.0282*<br>Interaction: F(3,6)= 3.879, p= 0.0742 | 24 h: 0 (n= 2), 1 (n= 2), 2.5 (n= 2), 5 (n= 2)<br>48 h: 0 (n= 2), 1 (n= 2), 2.5 (n= 1), 5 (n= 1) |
| 5a | Ketamine<br>One-way ANOVA:<br>Luciferase assay: F(7,156)= 1.654, p= 0.1243 | n= 27, 15, 22, 21, 22, 22, 17, 18 |
| 5b | 2R,6R-HNK<br>One-way ANOVA:<br>Luciferase assay: Kruskal-Wallis (7) = 5.784, p= 0.5652 | n= 15, 16, 16, 16, 16, 16, 12, 12 |
| 5c | 2S,6S-HNK<br>One-way ANOVA:<br>Luciferase assay: F(7,111)= 1.439, p= 0.1970 | n= 16, 16, 16, 16, 16, 16, 12, 11 |
| 5d | Rolipram<br>One-way ANOVA:<br>Luciferase assay: F(7,51)= 0.4432, p= 0.8702 | n= 8, 8, 8, 7, 8, 8, 6, 6 |
| 5e | Phenelzine<br>One-way ANOVA:<br>Luciferase assay: Kruskal-Wallis (7) = 14.62, p= 0.0412 | n= 8, 8, 8, 8, 7, 8, 5, 6 |
| 5f | Carbamazepine<br>One-way ANOVA:<br>Luciferase assay: F(6,35)= 2.325, p= 0.0540 | n= 6/group |
| 6a | Fluoxetine<br>Two-way ANOVA:<br>Treatment: F(1,98)= 894.2, p< 0.0001*<br>Mutant: F(2,98)= 1.481, p= 0.2324<br>Interaction: F(2,98)= 1.481, p= 0.2324 | WT (n= 16, 16)<br>N501Y (n= 18, 18)<br>K417N (n= 18, 18) |
| 6b | Fluoxetine<br>Two-way ANOVA:<br>Treatment: F(1,30)= 116.4, p< 0.0001*<br>Mutant: F(1,30)= 0.5339, p= 0.4707<br>Interaction: F(1,30)= 0.5339, p= 0.4707 | WT (n= 8, 8)<br>E484K (n= 9, 9) |
| 6c | Fluoxetine<br>Two-way ANOVA:<br>Treatment: F(1,54)= 255.5, p< 0.0001*<br>Mutant: F(1,54)= 0.4988, p= 0.4831<br>Interaction: F(1,54)= 0.4988, p= 0.4831 | WT (n= 16, 15)<br>Triple mutant (n= 14, 13) |
| 6d | WT-Fluoxetine<br>Two-way ANOVA:<br>Treatment: F(1,18)= 13.54, p= 0.0017*<br>Time: F(2,18)= 13.62, p= 0.0002*<br>Interaction: F(2,18)= 13.24, p= 0.0003* | n= 4/group |
| 6e | B.1.1.7-Fluoxetine<br>Two-way ANOVA:<br>Treatment: F(1,18)= 3.925, p= 0.0630<br>Time: F(2,18)= 3.930, p= 0.0384*<br>Interaction: F(2,18)= 3.811, p= 0.0417* | n= 4/group |
| 6f | B.1.351-Fluoxetine<br>Two-way ANOVA:<br>Treatment: F (1,18)= 484.1, p< 0.0001*<br>Time: F(2,18)= 463.2, p< 0.0001*<br>Interaction: F(2,18)= 459.1, p< 0.0001* | n= 4/group |
| 6, g, h, i | WT, B.1.1.7, B.1.351-Fluoxetine<br>Multiple ANOVA (MANOVA):<br>Treatment: F(1,18)= 9.810, p= 0.001<br>Genotype: F(2,18)= 1.746, p= 0.161<br>Interaction: F(2,18)= 1.397, p= 0,254 | n= 4/group |
| S2 | Camostat mesylate<br>One-way ANOVA:<br>Kruskal-Wallis (7) = 31.58, p< 0.0001* | n= 7, 7, 7, 7, 7, 7, 6, 6 |
| S3a | Fluoxetine<br>Two-way ANOVA:<br>Infection: F(1,8)= 1.893, p= 0.2062<br>Treatment: F(3,8)= 18.96, p= 0.0005*<br>Interaction: F(3,8)= 2.686, p= 0.1173 | n= 2/group |
| S3b | Citalopram<br>Two-way ANOVA:<br>Infection: F(1,8)= 23.45, p= 0.0013*<br>Treatment: F(3,8)= 0.9825, p= 0.4480<br>Interaction: F(3,8)= 0.8741, p= 0.4937 | n= 2/group |
| S3c | Venlafaxine | n = 3/group |

| | | |
|---|---|---|
| | Two-way ANOVA:<br>Infection: F(1,16)= 0.01145, p= 0.9161<br>Treatment: F(3,16)= 2.848, p= 0.0704<br>Interaction: F(3,16)= 1.977, p= 0.1581 | |
| S3d | Reboxetine<br>Two-way ANOVA:<br>Infection: F(1,16)= 2.801, p= 0.1137<br>Treatment: F (3, 16)= 4.212, p= 0.0225*<br>Interaction: F(3, 16)= 0.3931, p= 0.7597 | n= 3/group |
| S3e | Clomipramine<br>Two-way ANOVA:<br>Infection: F(1,8)= 4.762, p= 0.0607<br>Treatment: F(3,8)= 50.26, p< 0.0001*<br>Interaction: F(3,8)= 0.5625, p= 0.6547 | n= 2/group |
| S3f | Imipramine<br>Two-way ANOVA:<br>Infection: F(1,6)= 13.53, p= 0.0104*<br>Treatment: F(2,6)= 0.06564, p= 0.9371<br>Interaction: F(2,6) = 0.1169, p= 0.8916 | n= 2/group |
| S3g | Chlorpromazine<br>Two-way ANOVA:<br>Infection: F(1,8)= 57.15, p< 0.0001*<br>Treatment: F(3,8)= 1.342, p= 0.3277<br>Interaction: F(3,8)= 0.8214, p= 0.5178 | n= 2/group |
| S3h | Fluoxetine<br>Two-way ANOVA:<br>Interaction: F(1,12)= 0.1856, p= 0.6742<br>Treatment: F(1,12)= 0.3228, p= 0.5804<br>Virus: F (1,12)= 0.1857, p= 0.6742 | n= 4/group |
| S4a | Fluoxetine:<br>One-way ANOVA:<br>Luciferase: F(3,39)= 6.634, p= 0.0010*<br>Cell viability: F(4,19)= 3.920, p= 0.0174* | Luciferase: n= 15, 11, 11, 6<br>Cell viability: n= 8, 4, 4, 4 |
| S4b | Fluoxetine:<br>Two-way ANOVA:<br>Treatment: F(3,125)= 39.26, p< 0.0001*<br>Virus: F(1,125)= 56.79, p< 0.0001*<br>Interaction: F(3,125)= 8.490, p< 0.0001* | |
| S4c | Clomipramine<br>One-way ANOVA:<br>Luciferase: F(5,51)= 4.259, p= 0.0026*<br>Cell viability: F(3,16)= 9.611, p= 0.0007* | Luciferase: n= 17, 6, 6, 6, 11, 11<br>Cell viability: n= 8, 4, 4, 4 |
| S4d | Clomipramine<br>Two-way ANOVA:<br>Treatment: F(5,146)= 39.23, p< 0.0001*<br>Virus: F(1,146)= 57.05, p< 0.0001*<br>Interaction: F(5,146)= 16.59, p< 0.0001* | |
| S4e | Chlorpromazine<br>One-way ANOVA:<br>Luciferase: F(3,34)= 9.231, p= 0.0001*<br>Cell viability: F(2,13)= 10.42, p= 0.0020* | Luciferase: n= 11, 6, 10, 11<br>Cell viability: n= 8, 4, 4 |
| S4f | Chlorpromazine<br>Two-way ANOVA:<br>Treatment: F(3,86)= 38.78, p< 0.0001*<br>Virus: F(1,86)= 42.66, p< 0.0001*<br>Interaction: F(3,86)= 9.855, p< 0.0001* | |
| S4g | Flupenthixol<br>One-way ANOVA:<br>Luciferase: F(3,36)= 5.700, p= 0.0027*<br>Cell viability: F(3,16)= 0.3592, p= 0.7833 | Luciferase: n= 17, 6, 6, 11<br>Cell viability: n= 8, 4, 4, 4 |
| S4h | Flupenthixol<br>Two-way ANOVA:<br>Treatment: F(3,80)= 59.09, p< 0.0001*<br>Virus: F(1,80)= 66.26, p< 0.0001*<br>Interaction: F(3,80)= 12.42, p< 0.0001* | |
| S4i | Pimozide<br>One-way ANOVA:<br>Luciferase: F(2,27)= 5.234, p= 0.0120*<br>Cell viability: F(2,13)= 5.578, p= 0.0178* | Luciferase: n= 15, 5, 10<br>Cell viability: n= 8, 4, 4 |
| S4j | Pimozide<br>Two-way ANOVA:<br>Treatment: F(2,60)= 34.24, p< 0.0001*<br>Virus: F(1,60)= 4.542, p= 0.0372*<br>Interaction: F(2,60)= 2.295, p= 0.1096 | |
| S4k | Reboxetine | Luciferase: n= 11, 14, 14 |

| | | |
|---|---|---|
| | One-way ANOVA:<br>Luciferase: $F(2,36)= 4.344$, $p= 0.0204$*<br>Cell viability: $F(2,13)= 14.67$, $p= 0.0005$* | Cell viability: n= 8, 4, 4 |
| S4l | Reboxetine<br>Two-way ANOVA:<br>Treatment: $F(2,69)= 26.49$, $p< 0.0001$*<br>Virus: $F(1,69)= 16.47$, $p= 0.0001$*<br>Interaction: $F(2,69)= 5.190$, $p= 0.0079$* | |
| S5a | Citalopram<br>Unpaired t-test:<br>Luciferase: $t(20)= 1.612$, $p= 0.1227$<br>Cell viability: $t(10)= 1.017$, $p= 0.3333$ | Luciferase: n= 11/group<br>Cell viability: n= 8, 4 |
| S5b | Citalopram<br>Two-way ANOVA:<br>Treatment: $F(1,58)= 36.77$, $p< 0.0001$*<br>Virus: $F(1,58)= 12.95$, $p= 0.0007$*<br>Interaction: $F(1,58)= 12.95$, $p= 0.0007$* | |
| S5c | Desipramine<br>One-way ANOVA:<br>Luciferase: $F(2,30)= 1.396$, $p= 0.2633$<br>Cell viability: $F(2,13)= 3.591$, $p= 0.0573$ | Luciferase: n= 11/group<br>Cell viability: n= 8, 4, 4 |
| S5d | Desipramine<br>Two-way ANOVA:<br>Treatment: $F(2,63)= 24.24$, $p< 0.0001$*<br>Virus: $F(1,63)= 49.05$, $p< 0.0001$*<br>Interaction: $F(2,63)= 12.26$, $p< 0.0001$* | |
| S5e | Paroxetine<br>One-way ANOVA:<br>Luciferase: $F(2,30)= 2.296$, $p= 0.1181$<br>Cell viability: $F(2,13)= 0.2985$, $p= 0.7469$ | Luciferase: n= 11/group<br>Cell viability: n= 8, 4, 4 |
| S5f | Paroxetine<br>Two-way ANOVA:<br>Treatment: $F(2,63)= 28.49$, $p< 0.0001$*<br>Virus: $F(1,63)= 44.34$, $p< 0.0001$*<br>Interaction: $F(2,63)= 11.19$, $p< 0.0001$* | |
| S5g | Fluvoxamine<br>Unpaired t-test:<br>Luciferase: $t(20)= 0.3183$, $p= 0.7536$<br>Cell viability: $t(10)= 1.378$, $p= 0.1983$ | Luciferase: n= 11/group<br>Cell viability: n= 8, 4 |
| S5h | Fluvoxamine<br>Two-way ANOVA:<br>Treatment: $F(1,56)= 8.557$, $p= 0.0050$*<br>Virus: $F(1,56)= 10.78$, $p= 0.0018$*<br>Interaction: $F(1,56)= 10.78$, $p= 0.0018$* | |
